# The RNA helicase KREH2 directly binds the RESC1-RESC2 complex in trypanosomatid mitochondrial RNA editing

**DOI:** 10.64898/2026.10.02.754006

**Authors:** Rutuja Yelmar, Luciano Graciani Dolce, Laura Tengo, Bernardo Papini Gabiatti, Jane Corwin, Lars O’Hara, Cody Goode, Sunil K. Sharma, Tyler Rodshagen, Zachary Goodall, Jorge Cruz-Reyes, Suzanne M McDermott, Eva Kowalinski

## Abstract

Trypanosomatids are medically important single-celled eukaryotic protists with exceptional RNA biology. They display highly unusual RNA biogenesis pathways, notably mitochondrial RNA editing, an essential post-transcriptional process transforming cryptic primary transcripts into functional mRNAs. Large multiprotein assemblies coordinate the RNA editing process, in which small guide RNAs (gRNAs) anneal to the pre-mRNAs and serve as templates for uridine insertion or deletion. Kinetoplast RNA Editing Helicases (KREHs) are important components of the RNA editing process that modifies large portions of pre-mRNA via the sequential annealing and dissociation of multiple gRNAs. However, how KREH helicases interact with components of the editing machinery, and which subunits mediate these interactions, remain unresolved. Here, we examined an association between the DEAH/RHA RNA helicase KREH2 and the RESC1-RESC2 protein heterodimer using proximity labeling and co-immunoprecipitation coupled to quantitative mass spectrometry *in vivo*. We reconstituted a minimal trimeric KREH2 peptide-RESC1-RESC2 complex *in vitro* and characterized novel direct binding of a KREH2 peptide to RESC1 using cryo-electron microscopy. These results establish a mechanistic link between the RNA helicase KREH2 and the RESC1-RESC2 module at the 5′-end of gRNAs, providing new avenues for studying the orchestration of RNA remodeling in trypanosomatid mitochondrial RNA editing.

## Introduction

Kinetoplastids are unicellular protozoan parasites that cause vector-borne diseases in humans and animals, including trypanosomiasis and leishmaniasis (Stuart et al. 2008; McDermott et al. 2026; Pareyn et al. 2025; Lejon et al. 2025; Davidge et al. 2026). The mRNA biogenesis for several of the essential mitochondrial respiratory chain subunits in kinetoplastids requires RNA editing - the insertion or deletion of uridines in/from the transcribed RNA sequence. During pre-mRNA editing, genetic information from two distinct primary transcripts is combined. On one hand, a series of maxicircles (approximately 20–40 kb) of mitochondrial DNA encodes cryptic pre-mRNAs; these either lack uridine nucleotides or contain additional ones, rendering the transcripts nonsensical. On the other hand, thousands of small guide RNAs (gRNAs), typically 50–60 nucleotides long, are predominantly transcribed from DNA minicircles (about 1 kb) and direct mRNA editing (Stuart et al. 2005; McDermott et al. 2026; Blum and Simpson 1990). Each gRNA base-pairs with a target region on its cognate pre-mRNA through a specific 5′ anchor region (Aphasizheva et al. 2020; Stuart et al. 2005; Aphasizheva et al. 2014; Cooper et al. 2019, 2022). In its central region, the corrective gRNA forms a duplex with the pre-mRNA, and mismatches indicate target sites for pre-mRNA editing accomplished by a large protein machinery, which transforms the pre-mRNAs into transcripts encoding functional proteins. While some transcripts are minimally edited using few gRNAs, several mRNAs undergo multiple successive rounds of editing in a process termed pan-editing, guided by the sequential annealing of multiple gRNAs (Stuart et al. 2005; Aphasizheva et al. 2020; McDermott et al. 2026). The editing process requires extensive RNA remodeling to resolve RNA secondary structures, remove RNA-coating proteins, and dissociate gRNA:mRNA duplexes after editing - yet the mechanistic details remain largely unresolved (Read et al. 2016; McDermott et al. 2026).

The editing of each gRNA:mRNA mismatch is catalyzed by the <u>R</u>NA <u>E</u>diting <u>C</u>atalytic <u>C</u>omplexes (RECC), a spectrum of assemblies including a defined set of scaffolding and variable functional subunits, such as exo- and endonucleases, terminal uridylyltransferase (TUTase), and RNA ligases; these activities conduct pre-mRNA cleavage, insertion or removal of uridines, and ultimately re-ligate the mRNA strand (reviewed in (Aphasizheva et al. 2020; McDermott et al. 2026; Davidge et al. 2026; Stuart et al. 2005). Editing entails the coupling of RECC with additional modules of the catalytically inert <u>R</u>NA <u>E</u>diting <u>S</u>ubstrate Binding <u>C</u>omplex (RESC) subunits. RESC is a large, dynamic, potentially heterogeneous assembly of around 20 proteins functionally divided into a guide RNA binding module (formerly GRBC or MRB) and a module responsible for editing progression (formerly REMC or *Tb*RGG2 complex) (Hashimi et al. 2009; Weng et al. 2008; Ammerman et al. 2010; Aphasizheva et al. 2014; McAdams et al. 2019; Read et al. 2016). High-resolution structural work has begun to capture different RECC and RESC assemblies, reinforcing the view that the complex contains functional sub-modules that undergo regulated compositional rearrangements to bind pre-mRNA, select gRNAs, and provide a scaffold for RECC-catalyzed editing (Liu et al. 2023; Dolce et al. 2023; Liu et al. 2026). The RESC1 and RESC2 subunits form an obligate heterodimer that functions both autonomously and as a component of multimeric RESC assemblies. The RESC1-RESC2 heterodimer recognizes gRNAs based on their 5’-triphosphate end, binds this feature, and thereby stabilizes gRNA levels by preventing endonucleolytic degradation (Liu et al. 2023; Dolce et al. 2023; Weng et al. 2008); RESC1 and RESC2 are essential for *Trypanosoma brucei* cell viability and normal growth; their depletion results in a global gRNA depletion and reduction of edited mRNA levels (Weng et al. 2008; Hashimi et al. 2009). In addition to the RECC and RESC complexes, <u>K</u>inetoplast <u>R</u>NA <u>E</u>diting <u>H</u>elicases (KREHs) assist in RNA editing. These helicases exhibit ATP-dependent double-strand (ds)RNA unwinding activity *in vitro* with model mRNA-gRNA duplexes (Li et al. 2011; Kumar et al. 2019, 2016; Hernandez et al. 2010; Yelmar et al. 2026) and have been proposed to remodel gRNA and mRNA structures, displace RNA-binding proteins, and promote dissociation of the gRNA:mRNA duplex following editing, but additional studies are needed to understand the underlying molecular mechanisms of each helicase. The small DEAD-box helicase KREH1 (formerly Hel61 or REH1) seems to be required for editing of A6 mRNA and possibly other transcripts (Dubey et al. 2023; Li et al. 2011; Missel et al. 1997). It participates in local remodeling and facilitates editing progression, at least in insect procyclic form cells. KREH1 lacks known protein partners and associates transiently with other editing proteins in an RNA-dependent manner (Missel et al. 1995, 1997; Corell et al. 1996; Dubey et al. 2023; Panigrahi et al. 2003, 2006; Li et al. 2011; Wackowski et al. 2024; Hashimi et al. 2008; Hernandez et al. 2010; Yelmar et al. 2026).

The large multidomain DEAH-helicase KREH2 is essential for all RNA editing, modulating editing complex interactions, editing fidelity, and gRNA utilization (Meehan et al. 2023). KREH2 is also a key factor in the developmental regulation of editing in *T. brucei* (Hernandez et al. 2010; Madina et al. 2015, 2014; Meehan et al. 2023, 2024; Kumar et al. 2016; Hashimi et al. 2008). Native KREH2 ribonucleoprotein complexes (RNPs) contain protein cofactors, pre-mRNAs, partially edited RNAs, and mature edited products. These KREH2 RNPs retain mRNA but lose RESC association upon depletion of gRNAs via RESC2 depletion (Kumar et al. 2016). KREH2 crosslinks directly to editing sites in synthetic mRNA-gRNA duplexes or gRNAs *in vitro*, and exhibits 3′-5′ double-stranded RNA unwinding activity (Kumar et al. 2016; Hernandez et al. 2010; Kumar et al. 2019; Madina et al. 2015).

Topological models of KREH2 have been proposed (Cruz-Reyes et al. 2016; Kumar et al. 2016; McDermott et al. 2026), based on homology with the spliceosomal DEAH/RHA helicase Prp43 from yeast (Martin et al. 2002), and MLE that functions in Drosophila dosage compensation (Prabu et al. 2015; Jagtap et al. 2023). Early bioinformatic analyses identified two conserved double-stranded RNA-binding domains (dsRBDs). However, recent AlphaFold analyses predicted a total of six dsRBDs, including the original two within a long, flexible N-terminal region (residues 1-1153) (McDermott et al. 2026).

The C-terminal domain cluster (1154-2167) of KREH2 includes a dsRBD (dsRBD7), two RecA-like helicase domains, a helicase-associated domain 2 (HA2), and an oligonucleotide binding fold (OB-fold) (Cruz-Reyes et al. 2016; Kumar et al. 2016; McDermott et al. 2026). KREH2 forms a trimeric complex, termed REH2C, with <u>K</u>RE<u>H2</u>-associated <u>F</u>actors 1 and 2 (KH2F1 and KH2F2), as determined by reciprocal co-immunoprecipitation and mass spectrometry for each protein component (Hernandez et al. 2010; Kumar et al. 2016). Direct association of REH2C components was verified by purifications from mitochondrial extracts in the presence of RNase, from akinetoplastic bloodstream cells lacking mitochondrial RNA, and by reconstitution of recombinant protein complexes treated with RNase (Hernandez et al. 2010; Kumar et al. 2016; Meehan et al. 2024). KH2F1, a 58 kDa protein, consists of eight zinc-finger motifs of the Cys₂-His₂ (C2H2) subtype, is essential for cell growth and editing, and serves as an adaptor directing KREH2 to RESC and RECC (Kumar et al. 2016). KH2F1 RNAi-mediated knockdown reduces KREH2 association with RESC2, RESC13, KREL1, and pre-mRNA in co-immunoprecipitation experiments and decreases editing of the substrates examined *in vivo* (Kumar et al. 2016; Meehan et al. 2024). Deep sequencing demonstrated that KH2F1 knockdown alters REH2C-dependent gRNA utilization in multiple mRNAs during development (Kumar et al. 2020; Meehan et al. 2023, 2024). KH2F2 is a 30 kDa protein homologous to hydrolase domains. Its role is unclear, as its presence was not required for KREH2 association with KH2F1 or RESC, and its RNAi knockdown did not have apparent effects on editing (Kumar et al. 2016). Apart from KH2F1 and KH2F2, additional direct binding partners of KREH2 have not been reported.

Cooperation between the REH2C helicase complex and the catalytic RECC and scaffolding RESC has been substantiated by a range of approaches, including sedimentation analyses, co-immunoprecipitation, genetic studies, and deep sequencing of RESC-associated edited mRNA in REH2C mutants (Kumar et al. 2016; Madina et al. 2014, 2015; Hashimi et al. 2008; Meehan et al. 2023, 2024). Observations that KREH2 dissociated from RESC and RECC under stringent washing conditions or RNase treatment suggested that in some cases its association is transient and RNA-dependent (Hernandez et al. 2010; Liu et al. 2023). RECC and RESC complexes display substantial compositional diversity and functional specialization (McDermott et al. 2026). For example, canonical RESC but not RESC* contains the RESC subunits RESC5 and RESC6 (reviewed in (Cruz-Reyes et al. 2018; McDermott et al. 2026). KREH2 stably associates with RESC*, whereas RESC6 immunoprecipitations show minimal association with KREH2. Both RESC variants contain a RESC1-RESC2 heterodimer, suggesting that KREH2 associates with specific RESC configurations to fulfill distinct functional roles (Kumar et al. 2016). This is supported by changes in gRNA utilization, editing and abundances of mRNAs contained in RESC6 immunoprecipitates upon KREH2 knockdown (Madina et al. 2014, 2015; Kumar et al. 2016; Meehan et al. 2024, 2023).

Here, we dissect the proximity interactome of KREH2 using Turbo-ID in *Trypanosoma brucei* cells, and use co-immunoprecipitation of affinity-tagged KREH2 to examine interaction partners of KREH2. Our data indicate the trimeric REH2C complex with its two cofactors KH2F1 and KH2F2, as originally reported (Kumar et al. 2016). Additionally, the data identify a specific association between KREH2 and RESC1, and a proximal association between KH2F1 and RESC1. We further refine the domain topology model of KREH2 using AlphaFold and provide evidence for a direct interaction between KREH2 and the RESC1 subunit using *in vitro* interaction analysis and single-particle cryo-electron microscopy. Collectively, our findings corroborate the topology of the REH2C complex and establish a novel direct interaction between KREH2 and the RESC1–RESC2 heterodimer.

## Methods

### AlphaFold analysis

For individual protein analysis, prediction for KREH2 (Tb927.4.1500; uniprot: Q581T1), KH2F1 (Tb927.6.1680; uniprot: Q585P3) and KH2F2 (Tb927.6.2140; uniprot: Q584U2) (Aslett et al. 2010) were downloaded from AlphaFold database (Jumper et al. 2021; Varadi and Velankar 2023) under accession numbers AF-Q581T1-F1-v6, AF-Q585P3-F1-v6, and AF-Q584U2-F1-v6, respectively. Domain boundaries were defined by manual inspection of predicted aligned error (PAE) matrices, and were annotated based on sequence and/or structural similarity. To predict protein-protein interaction, the KREH2 sequence was divided into 5 fragments corresponding to the defined domains, and each fragment was modeled in complex with the cofactors and RESC components using AlphaFold-Multimer through ColabFold (Jumper et al. 2021; Mirdita et al. 2022). Predictions with low intermolecular PAE scores were repeated using AlphaFold3 (Abramson et al. 2024) (available online at alphafoldserver.com), focused on the interaction regions to refine the results.

### *Trypanosoma brucei* cell culture and cell-line generation

Procyclic form SmOx cells (Poon et al. 2012; Delauw et al. 1985) and AnTat 1.1 *T. brucei* cells (Poon et al. 2012; Delauw et al. 1985) were cultured in complete SDM-79 medium supplemented with 10% heat-inactivated fetal bovine serum, 20 μg/ml penicillin-streptomycin, 0.4% glycerol, and 30 μg/ml hemin at 27°C (Brun and Schönenberger 1979). Cells were maintained in a logarithmic growth phase at 1 × 10^6^ cells/ml by subculturing every second day. Before experiments, cells were subcultured at least twice after thawing.

To generate cell lines for proximity labeling experiments, the KREH2 and KH2F1 genes were endogenously tagged using a PCR-based tagging strategy (Dean et al. 2015; Billington et al. 2023) with a modified pPOT vector containing TurboID-HA (Susanne Kramer, University of Würzburg, Germany) used as the PCR template. Briefly, PCR products were ethanol-precipitated by mixing 0.1 volumes of 3 M sodium acetate with 3 volumes of 100% EtOH, incubating for 1 h at -80°C, and washing twice with 70% EtOH. A total of 10 µg of purified PCR product was transfected into 2 × 10^7^ cells in 400 µl of cytomix buffer (25 mM HEPES pH 7.6, 120 mM KCl, 0.15 mM CaCl2, 10 mM K2HPO4/KH2PO4 pH 7.6, 2 mM EDTA, 6 mM glucose, 5 mM MgCl2, sterile filtered) using a BTX Gemini SC2 electroporator (800 V, 50 µF, 100 Ω). Transfected cells were resuspended in 22.5 ml SDM-79 medium supplemented with 2.5 ml FBS (final 20% FBS) and incubated at 27°C with 5% CO₂ for 18 h to allow recovery. After recovery, 13 ml of the recovered culture was diluted with 34 ml of complete medium, 4 ml of FBS, and 20 µg/ml blasticidin and plated into two 24-well plates (1 ml/well). Plates were sealed and incubated at 27°C with 5% CO₂ for 10-11 days until cultures were sufficiently dense for expansion of positive clones in 10 ml of medium containing 10 µg/ml blasticidin. A control cell line was generated similarly, except that the N-terminal mitochondrial targeting sequence (MTS) from KREH1 (predicted with MitoFates) was fused to enhanced YFP-TurboID-HA (referred as MTS-eYFP henceforth) in the tubulin locus for constitutive expression (Fukasawa et al. 2015; Moreira et al. 2023).

For the generation of cell lines for co-immunoprecipitation experiments, the C-terminal tagging cassette comprising the Streptavidin-binding peptide (SBP)-3C cleavage site (3C)-Protein A (ProtA) tag and neomycin resistance was PCR-amplified from a modified pPOT vector using KREH2 and RESC2 (Tb927.2.3800) endogenous tagging primers (Dean et al. 2015; Billington et al. 2023) and transfected into AnTat 1.1 cells. For selection, transfected cells were transferred to 5 ml of antibiotic-free conditioned medium and incubated for 24 h before the addition of 40 µg/ml neomycin for selection. After another 7 days, the neomycin concentration was reduced to 20 µg/ml. Recovery of positive clones was achieved after approximately 3 weeks. All transgenic cell lines generated in this study are indicated in Supplementary Table S1.

### Generation of a genetic complementation system for KREH2 variants in bloodstream form cells

Bloodstream form (BSF) *T. brucei* strain Lister 427 Single Marker (SM) cells were cultured as described previously (Meehan et al. 2023, 2024). A single knockout (SKO) of KREH2 was introduced using standard techniques with modifications (McDermott et al. 2019, 2015). Briefly, we constructed a knockout vector containing a marker cassette, including blasticidin resistance, flanked by 5’ and 3’ UTR sequences of KREH2. The fragments were assembled in a three-piece assembly using an In-Fusion kit (Takara Bio) and then fused into the linearized vector backbone using the same kit. The complete marker cassette with flanking homology arms was amplified from the sequence-verified KO vector and transfected into trypanosomes. Correct cassette insertion was confirmed by PCR and Sanger sequencing (see Supplementary Tables S2 for primers and S3 for plasmids) and drug sensitivity assays. A tetracycline (tet)-inducible RNA interference (RNAi) construct targeting the endogenous 3′ untranslated region (UTR) (Meehan et al. 2023) was introduced into our SKO KREH2 cell line. Induction with 1 µg/mL tet (Sigma) reduced KREH2 protein by ∼80% by day 1, and difference in parasite growth began by day 2 in clonal lines of KREH2 SKO/RNAi cells obtained by limiting dilution. For genetic complementation, the 6,504 bp wild-type (WT) KREH2 coding sequence (CDS) bearing a stop codon was PCR-amplified and cloned by In-Fusion (Takara Bio) into a slightly modified version of pHD1344tub(PAC)GWCterm3V5 (McDermott et al. 2015), which uses a heterologous 3′ UTR. This vector renders the ectopic KREH2 allele RNAi-resistant and drives constitutive expression from the β-tubulin locus. The resulting pHD1344tub(PAC)-KREH2 WT plasmid was digested with NotI and used to transfect KREH2 SKO/RNAi cells. Transgenic lines were selected with puromycin, construct insertion was verified by PCR (Supplementary Table S2), and constitutive expression of untagged KREH2 was confirmed by Western blotting using anti-KREH2 and anti-eF1a antibodies (Meehan et al. 2023, 2024) as a loading control. To generate the KREH2 K564E/R566E/K569E/R570E variant, the pHD1344tub(PAC)-KREH2 WT plasmid was digested with AvrII and BsrGI to produce the necessary vector. A synthetic gene fragment (GenScript) containing the K564E/R566E/K569E/R570E mutation was amplified with primers containing AvrII and BsrGI sites (Supplementary Table S4). After digestion, the gene fragment was ligated into the vector via Gibson Assembly (NEBuilder HiFi DNA Assembly Cloning Kit, New England BioLabs). The resulting construct pHD1344tub(PAC)-KREH2 (K564E/R566E/K569E/R570E) was digested with NotI before transfection into BSF KREH2 SKO/RNAi cells. Transgenic clonal cell lines were genotype verified via Sanger sequencing. All plasmids were deep-sequenced (Supplementary Table S3, Plasmidsaurus).

For growth curve analysis, cell density was measured using a Coulter Counter. BSF cells growing in the absence vs presence of 1 µg/ml tetracycline were reseeded at 1 to 2 × 10^5^ cells/mL in 10 ml every day. To generate cell growth heat maps, cumulative growth numbers for +tet and -tet cultures were calculated for each time point. Next, log2 of the ratio of +tet to -tet was calculated. This ratio was converted to blue-orange scale using RColorBrewer (Neuwirth 2002). The heat map was generated in R (R Core Team 2026) using the gplots (Warnes et al. 2005), RColorBrewer (Neuwirth 2002), and readxl (Wickham and Bryan 2015) packages.

### Streptavidin affinity purification

Affinity purifications were carried out in three technical replicates following published protocols (Moreira et al. 2023; Odenwald et al. 2024). Endogenous biotin levels in SDM79 medium were sufficient for efficient TurboID-mediated labeling (Moreira et al. 2023). For each replicate, 5 × 10^8^ cells were harvested by centrifugation (1,400 × g, 10 min, 4°C), washed twice with serum- and hemin-free SDM79 medium, transferred to low-binding tubes (Eppendorf), snap-frozen, and stored at −80°C. Pellets were lysed in lysis buffer (50 mM Tris–HCl pH 7.5, 150 mM NaCl, 0.4% SDS, 1% IGEPAL CA-630, 1 mM EGTA, 1.5 mM MgCl₂, 0.25 mM DTT, and EDTA-free protease inhibitors (Roche)), incubated for 10 min at 4°C on a rotating wheel, and clarified by centrifugation (14,000 × g, 5 min). The clarified lysate was incubated with 50 µl of pre-equilibrated streptavidin magnetic beads (MagReSyn®, ReSyn Biosciences) for 1 h at 4°C. Beads were washed sequentially with lysis buffer (6×), 4 M urea in 50 mM TEAB pH 8.5 (2×), 6 M urea in 50 mM TEAB pH 8.5 (2×), 1 M KCl in 50 mM TEAB pH 8.5 (2×), and 50 mM TEAB pH 8.5 (2×) with each wash performed for 8 min at 4°C. On-bead digestion was performed by incubating beads in 40 µl of digestion buffer (50 mM TEAB pH 8.5, 10 mM DTT) in a thermomixer at 1000 rpm (1 h, RT), followed by alkylation with 20 mM iodoacetamide (30 min, in the dark, without shaking). Proteins were digested on-bead overnight at 30°C with 500 ng trypsin/LysC mix (Mass Spec Grade; Promega) in the presence of 1 mM biotin in a thermomixer at 1000 rpm. Supernatants were collected, and beads were subjected to a second on-bead digestion by incubation in 40 µl of digestion buffer containing 500 ng trypsin/Lys-C and 1 mM biotin at 30°C for 1 h in a thermomixer at 1000 rpm. Both supernatants were combined and dried using a SpeedVac (Eppendorf) at 30 ℃. The peptides were stored at -80 ℃ until mass spectrometry analysis.

### Mass spectrometry analysis of streptavidin purifications

Peptides were desalted on an OASIS® HLB µElution Plate (Waters), dried by vacuum centrifugation, and reconstituted in 1% formic acid, 4% acetonitrile. LC–MS analysis was performed on an UltiMate 3000 RSLCnano system (Thermo Fisher Scientific) equipped with a µ-Precolumn C18 PepMap™ 100 trapping cartridge (300 µM i.d. × 5 mm, 5 µM, 100 Å) and a nanoEase™ M/Z HSS T3 analytical column (75 µM i.d. × 250 mm, 1.8 µM, 100 Å; Waters). Samples were trapped at 30 µl/min using 0.05% trifluoroacetic acid (TFA) in water for 6 min and separated at 0.3 µl/min using solvent A (3% dimethyl sulfoxide (DMSO), 0.1% formic acid in water) and solvent B (3% DMSO, 0.1% formic acid in acetonitrile) with the following gradient: 2–8% in 6 min, 8–25% in 39 min, 25-40% in 5 min, 40–85% in 0.1 min, held at 85% for 3.9 min, and re-equilibrated to 2% for 6 min. The column outlet was directly coupled to an Orbitrap Fusion™ Lumos™ Tribrid™ mass spectrometer (Thermo Fisher Scientific) equipped with a Nanospray Flex™ ion source operating in positive-ion mode. Peptides were introduced using a Pico-Tip emitter (360 µM OD × 20 µM ID; 10 µM tip, CoAnn Technologies) at 2.4 kV, with a capillary temperature of 275°C. MS1 scans were acquired in centroid mode over m/z 420–680 at 60,000 resolution (at m/z 200), with AGC 300% and injection time set to ‘auto’. DIA was performed over m/z 430–670 using 12 m/z isolation windows. MS/MS scans were acquired in centroid mode over m/z 200–1800 at 30,000 resolution, with AGC 3000%, injection time set to ‘auto’, and higher-energy collisional dissociation (HCD) at 30%.

Raw files were analyzed using DIA-NN 1.8.1 (Demichev et al. 2020) in directDIA mode with an in silico-generated spectral library from *Trypanosoma brucei* FASTA files (TREU927_release67; 21 Feb 2024) (Aslett et al. 2010). Search parameters included Trypsin as protease; C carbamidomethylation and N-terminal M excision as fixed modifications; Protein N-terminal acetylation and Methionine oxidation as variable modifications; 2 missed cleavages; precursor m/z range 430–670; and charge states 2–4. DIA-NN settings were: Precursor FDR (%) = 1; Mass accuracy, MS1 accuracy, and Scan window = 0; isotopologues enabled; match between runs enabled; no shared spectra; protein inference at the gene level; single-pass neural network classifier; Robust LC (high precision) quantification; RT-dependent cross-run normalization; Smart profiling library generation; and optimal speed/RAM usage. Protein-level results were filtered using Q.Value ≤ 0.01, PG.Q.Value ≤ 0.01, and Lib.PG.Q.Value ≤ 0.01, and quantified using the dia-nn R package implementing DIA-NN’s MaxLFQ algorithm (Demichev et al. 2020). The final output (dia-nn_outputs_filtered_report.uniqueGenes_maxLFQ_matrix.tsv) contained gene-level intensities with Razor, Unique, and Total peptide counts, and Total Intensity is available as Supplementary Table S5. Differential protein abundance was assessed using limma. Proteins were classified as hits at FDR ≤ 0.05 and absolute fold change ≥ 1.5, and as candidates at FDR ≤ 0.2 and absolute fold change ≥ 1.5. Positive-fold-change hits and candidates in the corresponding tagged-construct versus YFP comparisons were classified as enriched hits and enriched candidates, respectively. Results were visualized as volcano plots with GraphPad Prism. The mass spectrometry proteomics data have been deposited to the ProteomeXchange Consortium via the PRIDE partner repository with the dataset identifier PXD081786 (Perez-Riverol et al. 2022).

### Co-immunoprecipitation of KREH2 and RESC2

2 × 10^10^ cells were harvested, washed once with 20 ml 1× PBS, and resuspended in 10 ml of immunoprecipitation (IP) buffer (200 mM NaCl, 10 mM MgCl2, 20 mM HEPES, 1 mM DTT, and a Complete protease inhibitor tablet (Roche)). Cells were lysed by sonication with a 5-s on, 15-s off cycle for a total of 4 min at 30% amplitude on ice (Sonics VC750 sonicator). The lysate was cleared by centrifugation at 12,096 × g for 60 min at 4 ℃, and the supernatant was divided into three technical replicates, each containing approximately 180 mg of protein, with protein concentration estimated by absorbance at 280 nm using a NanoDrop spectrophotometer. Each replicate was incubated with 100 µl of IgG sepharose resin (Cytiva), pre-equilibrated with IP buffer, for 1 h at 4 ℃ in an orbital shaker. Resin was spun down at 900 × g for 1 min at 4 ℃, and the supernatant was discarded. Resin was washed twice with 500 µl of IP buffer (without protease inhibitors), each time followed by centrifugation at 900 × g for 1 min at 4 ℃. For samples treated with RNase, the lysate mixed with resin was subsequently incubated with 20 µl of RNase A/T1 mix (2 mg/ml stock; Thermo Scientific^TM^) in 500 µl of IP buffer for 1 h at 4 ℃; untreated samples were incubated under identical conditions in IP buffer without RNase. After this step, the resins were washed twice with 500 µl of IP buffer (without protease inhibitors), each time followed by centrifugation at 900 × g for 1 min at 4 ℃. To elute the bound proteins, the resins were incubated with 500 µl of IP buffer containing 10 µl of 3C protease (1 mg/ml; PEPCF, EMBL Heidelberg) in an orbital shaker for 1 h at 4 ℃ and 50 µl of the eluate was collected separately from this step and the remaining 450 µl of eluate was bound to 50 µl pre-equilibrated Streptavidin-agarose resins (Thermo Scientific^TM^) and incubated overnight at 4 ℃ on a rotating wheel. The SBP-unbound fraction was removed by centrifugation at 900 × g for 1 min at 4 ℃. The resins were washed twice with 500 µl of IP buffer. The SBP-bound protein was eluted by incubating the resins with 100 µl of IP buffer containing 10 mM desthiobiotin (IBA) in a thermomixer at 800 rpm for 2 h at 4 ℃, followed by collection of the supernatant containing soluble protein. Eluates from both steps were analyzed using multiplex mass spectrometry.

### Mass spectrometry analysis of co-immunoprecipitations

Disulfide bonds were reduced with dithiothreitol (10 mM in 50 mM HEPES, pH 8.5; 56 °C, 30 min) and alkylated with 2-chloroacetamide (20 mM in 50 mM HEPES, pH 8.5; room temperature, dark, 30 min). Samples were cleaned using the SP3 protocol (Hughes et al. 2014, 2019) and digested overnight at 37 °C with sequencing-grade trypsin (Promega) at a 1:50 enzyme-to-protein ratio. Peptides were labeled with TMT10plex (Thermo Fisher; (Werner et al. 2014) according to the manufacturer’s instructions: 0.8 mg of reagent in 42 µl of acetonitrile (100%), 8 µl added per sample, incubated for 1 h at room temperature, and quenched with 8 µl of 5% hydroxylamine for 15 min. Labeled samples were combined, desalted on an OASIS® HLB µElution Plate (Waters), dried by vacuum centrifugation, reconstituted in 30 µl 1% formic acid, 4% acetonitrile, and stored at −80 °C until analysis.

Peptides were analyzed on an UltiMate 3000 RSLCnano system (Thermo Fisher Scientific) with a µ-Precolumn C18 PepMap™ 100 trapping cartridge (300 µM i.d. × 5 mm, 5 µM, 100 Å) and a nanoEase™ M/Z HSS T3 analytical column (75 µM i.d. × 250 mm, 1.8 µM, 100 Å; Waters). Samples were trapped at 30 µl/min in 0.05% trifluoroacetic acid (TFA) for 6 min and separated at 0.3 µl/min using solvent A (3% DMSO, 0.1% formic acid in water) and solvent B (3% DMSO, 0.1% formic acid in acetonitrile) with the gradient: 2–8% in 6 min, 8–25% in 39 min, 25–40% in 5 min, 40–85% in 0.1 min, held at 85% for 3.9 min, and re-equilibrated to 2% for 6 min. Peptides were introduced into a Q Exactive™ Plus mass spectrometer (Thermo Fisher Scientific) via a Pico-Tip emitter (360 µM OD × 20 µM ID; 10 µM tip, CoAnn Technologies) at 2.2 kV, with capillary temperature 275°C. MS1 scans were acquired in profile mode over m/z 375–1,200 at 70,000 resolution (at m/z 200), injection time 250 ms, and AGC 3E6. DDA MS/MS scans were acquired at 35,000 resolution, injection time 120 ms, AGC 2E5, HCD 30%, quadrupole isolation 0.7 m/z, dynamic exclusion 30 s, peptide match ‘preferred’, charge exclusion ‘unassigned’, selecting precursors 2–4+. Raw files were converted to mzml (MSConvert, ProteoWizard) with peak picking, 64-bit encoding, zlib compression, and top 1000 peaks retained. Data were searched with MSFragger (FragPipe 22.1-build02) against the *Trypanosoma brucei* FASTA (TREU927_release67; 21 Feb 2024; 9788 proteins) (Aslett et al. 2010), including contaminants and decoys. Fixed modifications: Carbamidomethylation (C, 57.0215) and TMT10 (K, 229.1629); variable: Oxidation (M, 15.9949) and TMT10 (peptide N-term, 229.1629). Trypsin specificity (≤2 missed cleavages, peptide length ≥7) and 20 ppm MS1/MS2 tolerances were used. Peptide/protein false discovery rate (FDR) was controlled at 1%. Quantitative analysis was performed in R (ISBN 3-900051-07-0). Contaminants were removed, retaining 1,318 proteins with ≥2 razor peptides. Replicate batch effects were corrected with removeBatchEffect (Ritchie et al. 2015) followed by variance stabilization normalization with normalizeVSN (Huber et al. 2002). Differential expression analysis was performed using the moderated t-test implemented in limma, with replicate information included as a factor in the design matrix (Ritchie et al. 2015). The fdrtool function was used to analyze the t-values produced by limma for the KREH2 versus AnTat 1.1 and RESC2 versus AnTat 1.1 comparisons within each sample type (RNase treated, RNase untreated) (Strimmer 2008). For the comparisons, P-values were adjusted for multiple testing using the Benjamini–Hochberg procedure. Proteins were annotated as hits at false discovery rate (FDR) ≤ 0.05 and |Fold Change| ≥1.5; enriched candidates were defined at FDR ≤ 0.2 and |Fold Change| ≥1.5. Results were visualized as volcano plots (GraphPad Prism). The mass spectrometry proteomics data have been deposited to the ProteomeXchange Consortium via the PRIDE partner repository with the dataset identifier PXD081786 (Perez-Riverol et al. 2022).

### Protein Expression and Purification

RESC1–RESC2 heterodimer was purified with an established protocol (Dolce et al. 2023). RESC1 (residues 22–473) carrying an N-terminal 6×His-3C tag and untagged RESC2 (residues 33–492), both lacking their predicted mitochondrial targeting sequences, were co-expressed in Hi5 insect cells using the BigBac system. Cell lysates were subjected to Ni²⁺-affinity chromatography, followed by tag removal with 3C protease and further purification by anion-exchange chromatography. The complex was finally polished by Superdex 200 Increase 10/300 GL size-exclusion chromatography in 20 mM HEPES pH 7.5, 150 mM NaCl, 2 mM MgCl₂, 1 mM DTT. All purification steps were performed at 4 °C.

### Isothermal Titration Calorimetry

KREH2 peptides were synthesized by SB-Peptide (Saint-Égrève, France) as wild-type (PQNT**KRR**VA**KR**ARVADAQPT), single inverse-charge mutants R565E (PQNTK**E**RVAKRARVADAQPT), R566E (PQNTKR**E**VAKRARVADAQPT), K569E (PQNTKRRVA**E**RARVADAQPT), R570E (PQNTKRRVAK**E**ARVADAQPT), and a quadruple inverse-charge mutant peptide (PQNT**E**R**E**VA**EE**ARVADAQPT). The peptides and the RESC1-RESC2 complex were dialyzed against ITC buffer containing 20 mM HEPES (pH 7.5), 50 mM NaCl, 2 mM MgCl₂, and 1 mM DTT. ITC experiments were performed at 20 °C using a MicroCal PEAQ-ITC calorimeter (Malvern). The peptide at 200 µM was titrated from the injection syringe into the sample cell containing 20 µM protein. Control titrations of peptide into buffer alone were carried out under identical conditions, and the resulting heats were subtracted from sample titrations to correct for heats of dilution. Raw thermograms were integrated using MicroCal PEAQ-ITC Analysis Software (Malvern) to obtain normalized heats per injection. Normalized heats were plotted as a function of molar ratio. Each titration experiment was fitted individually to a one-set-of-sites binding model using MicroCal PEAQ-ITC Analysis Software. Binding parameters (dissociation constant Kd, binding enthalpy ΔH, and stoichiometry n) were obtained from the fits. Reported values represent the mean ± standard deviation. Three independent titration experiments were performed for the wild-type peptide, two for the single mutants with reduced binding, and a single experiment for the peptides that did not exhibit binding.

### Cryo-EM sample preparation and data acquisition

For Cryo-EM sample preparation, purified RESC1-RESC2 (0.5 μM) was mixed with wild-type KREH2 peptide (5 μM) in 80 µl ITC buffer (20 mM HEPES pH 7.5; 50 mM NaCl; 2 mM MgCl2; 1 mM DTT) and incubated on ice for 30 min. After incubation, 2.5 µl of the sample was applied to both sides of UltrAuFoil R 1.2/1.3 300 Au mesh grids (Quantifoil), which were previously plasma-cleaned for 90 s using a 75:25 (Ar:O2) gas mixture. Excess sample was blotted with 0 blot force for 3 s and vitrified in liquid ethane using a Vitrobot MARK IV (Thermo Fisher Scientific) operated at 4°C and 100% humidity.

Movies were collected with a 300 kV Titan Krios (Thermo Fisher Scientific) electron microscope at ESRF (CM02), equipped with a Cold FEG, a Falcon4i direct electron detector, and a SelectrisX energy filter operated with a slit width of 10 eV. Data was acquired in an automated manner using EPU at a nominal magnification of 165,000x, resulting in a pixel size of 0.73 Å. To improve sampling of particle orientations, the stage was tilted 30°. Movies were acquired for 7.34 s at a dose of 5.11 e/pixel/s, resulting in a total sample dose of 64.95 e⁻/Å², fractionated into 126 movie fractions. Four movies were acquired per hole, totaling 7500 movies, with a defocus range of −1.2 to −2.6 μm.

### Cryo-EM data processing

Cryo-EM data were processed in CryoSPARC (v4.7.1, Structural Biotechnology Inc). Movies were motion corrected via Patch Motion Correction, and CTF was estimated using Patch CTF estimation. Micrographs were curated based on global motion, defocus range, estimated resolution, and ice thickness. The remaining micrographs were denoised and randomly split into subsets of 1000 micrographs. One subset was used for particle picking with Blob Picker, using an elliptical blob. Picked particles were curated automatically with Junk Detector and Inspect Particle Picks in Clustering mode (CryoSPARC Team et al. 2025), extracted, and 2D classified. Good classes were used for template pick in the full dataset. A total of 706,131 particles (400-pixel box) were extracted, binned, and used to generate 3 ab initio reconstructions, followed by heterogeneous refinement. Particles of a peptide-bound class were used for a second round of ab initio and heterogeneous refinement with 3 classes, from where the particles of the 2 good classes were pooled together. The final set of 273,716 particles was used for non-uniform refinement, yielding a refined map with an estimated resolution of 2.9 Å, as calculated by the CryoSPARC gold-standard FSC method. Final map was postprocessed with DeepEMhancer (Sanchez-Garcia et al. 2021).

### Model building

The model was generated based on our previous structure of RESC1-RESC2 heterodimer (Dolce et al. 2023) (PDB: 8CDP). The previous RESC1-RESC2 model was fitted to the map using ChimeraX (Meng et al. 2023) and refined against the nonsharpened map interactively, with Ramachandran and secondary-structure restraints, using both Coot 0.9.8 (Casañal et al. 2020) and Phenix (Liebschner et al. 2019). The peptide was generated manually in Coot 0.9.8.

### RNA isolation, and RT-PCR analyses of RNA editing

Total RNA was harvested using TRIzol and treated with TURBO DNase (Thermo Fisher Scientific) according to manufacturer’s instructions from cells grown 24 h after RNAi induction via tet addition. RNA integrity was confirmed using an RNA nanochip on a BioAnalyzer (Agilent Technologies). For high-throughput real-time PCR of endogenous vs. constitutively expressed WT and mutant KREH2, and never-, pre- and fully-edited mitochondrial mRNAs on the BioMark HD system (Standard BioTools), total RNA was reverse transcribed using TaqMan Reverse Transcription Reagents and MultiScribe Reverse Transcriptase (Thermo Fisher Scientific), pre-amplified in multiplex Specific Target Amplification (STA) reactions using TaqMan PreAmp Master Mix (Thermo Fisher Scientific) and treated with Exonuclease I (NEB). The abundances of reference telomerase reverse transcriptase (TERT), KREH2, and never-, pre- and fully-edited transcript cDNAs were then analyzed as previously described (McDermott et al. 2015). Primers are described in Supplementary Table S6. Calculations of RNA levels in samples following 24 h tet addition, relative to the absence of tet, were done using the 2 [-ΔΔC(T)] method (Livak and Schmittgen 2001). At least two technical duplicates of each cDNA sample were assayed for each target and internal reference per experiment and C(T) data averaged before performing the 2 [−ΔΔC(T)] calculation. Experiments were repeated using at least two biological replicates.

## Results

### The REH2C complex topology based on AlphaFold

According to the AlphaFold protein structure database model, the KREH2 catalytic DEIH motif is embedded within a globular C-terminal domain (residues 1154–2140) (AF-Q581T1-F1-v6, Figure 1A, B), which is in agreement with reported homology and more recent AlphaFold3 models of KREH2 (Kumar et al. 2016; Jumper et al. 2021; Abramson et al. 2024; McDermott et al. 2026). In addition to the two RecA domains, this domain comprises a double-stranded RNA-binding domain (dsRBD), a helicase-associated domain 2 (HA2) and an OB-fold-like domain. Similar helicase clusters are characteristic of DEAH/RHA RNA helicases, such as Prp2, Prp16, Prp22 and Prp43 splicing helicases in *Saccharomyces cerevisiae,* and their homologs, human DHX9 or *Drosophila melanogaster* Maleless (MLE), a core component of the dosage compensation complex (DCC) (Prabu et al. 2015; Jagtap et al. 2023; Tauchert et al. 2017; Walbott et al. 2010; He et al. 2010; Wilkinson et al. 2017; Cruz-Reyes et al. 2016; McDermott et al. 2026; Cristini et al. 2018). Compared with Prp43 (Tauchert et al. 2017), the residues that define the RNA-binding tunnel are conserved, suggesting a similar mode of RNA binding (Figure S1). The N-terminal half of KREH2 (residues 1-1153) contains six individual dsRBDs as was previously reported (McDermott et al. 2026), lacking interdomain correlation reminiscent of beads on a string; the seventh dsRBD is associated with the helicase domain cluster (Figure 1A, B, S2A). All KREH2 dsRBDs include an extra C-terminal α-helix compared to the canonical fold; dsRBD3, dsRBD4, dsRBD5, and dsRBD6 feature additional insertions (Figure S2B, C). dsRBD1, dsRBD5 and dsRBD6 contain positively charged surface patches, that are potentially able to interact with nucleic acids, suggesting that the N-terminus of KREH2 contributes to RNA binding. Such a role is in line with previous *in vivo* studies of KREH2 dsRBD mutants, as well as recent structural models (Hernandez et al. 2010; Cruz-Reyes et al. 2016; McDermott et al. 2026; Kumar et al. 2019) (Figure S2C).

**Figure 1:**
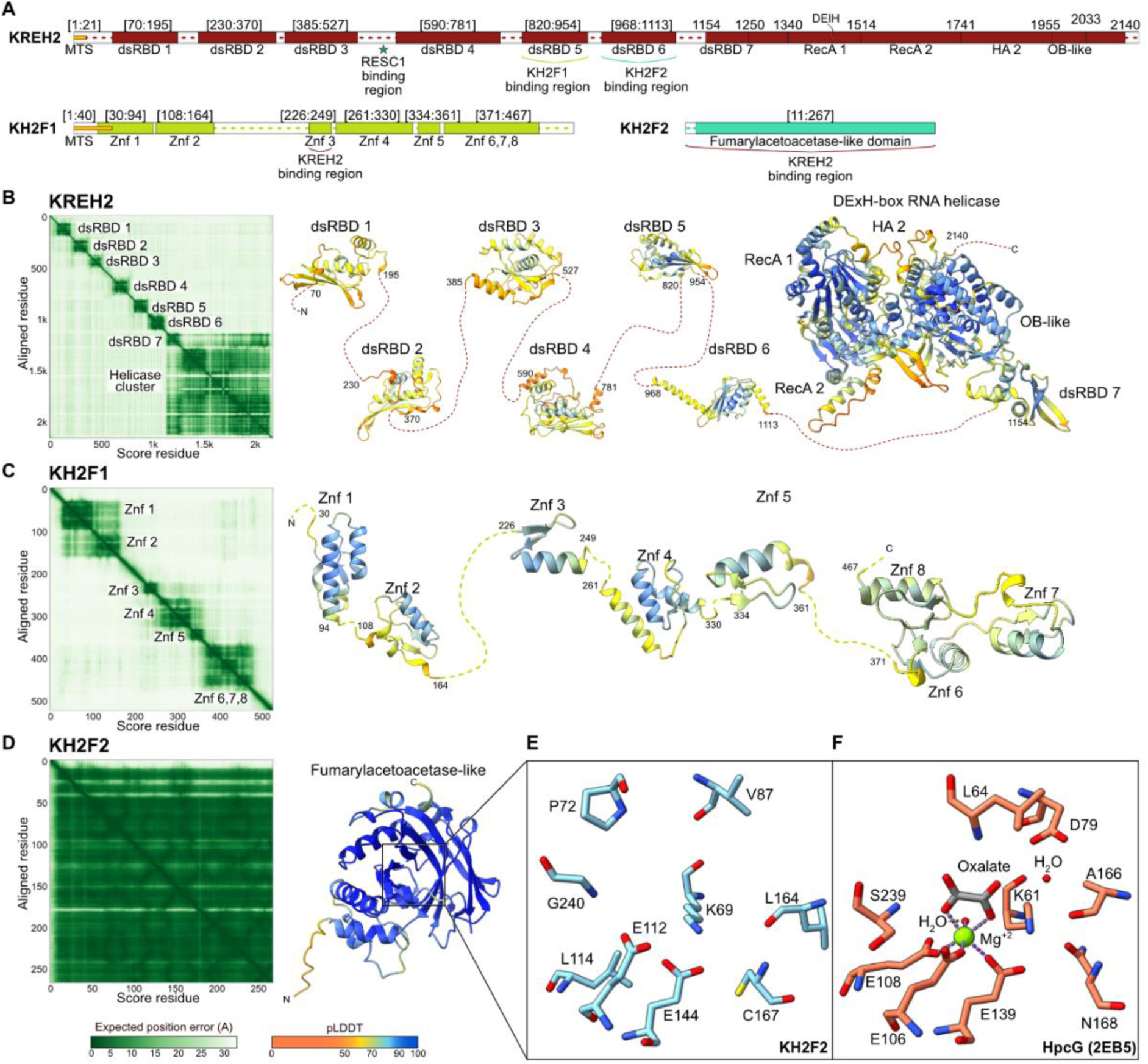
AlphaFold models of the REH2C complex components. (**A**) Schematic of KREH2 (dark red), KH2F1 (light green), and KH2F2 (light blue), indicating the domain organization based on AlphaFold models. The mitochondrial targeting sequence is shown in yellow, and disordered regions are indicated by dashed lines. (**B**) KREH2 (AF-Q581T1-F1-v6) AlphaFold model. (**C**) KH2F1 (AF-Q585P3-F1-v6) AlphaFold model. (**D**) KH2F2 (AF-Q584U2-F1-v6) AlphaFold model. In (B-D), the Predicted Aligned Error (PAE) plot, and the cartoon colored by pLDDT score are shown; dashed lines represent disordered regions. (**E**) Zoomed-in view of the active-site–homologous region of KH2F2. (**F**) in comparison with the substrate-bound active site of HpcG (2EB5). Stick representation of side chains, water molecules as red spheres, oxalate in grey, and Mg^2+^ as a green sphere with dotted lines indicating its coordination contacts.

The first helicase cofactor KH2F1 contains eight Cys₂-His₂ (C2H2) zinc finger (Znf) domains (AF-Q585P3-F1-v6, Figure 1A-C, S3A-C) (Kumar et al. 2016), organized in three independent clusters - Znf1 and Znf2; Znf3, Znf4 and Znf5; and Znf6, Znf7 and Znf8 (Figure 1C); similar Znf ensembles can function cooperatively as modular arrays to mediate nucleic acid binding (Guo et al. 2018; Yang et al. 2023; Fedotova et al. 2017). AlphaFold3 predicts that Znf3 of KH2F1 interacts with a positively charged surface patch on dsRBD5 of KREH2, which we confirmed by a relocalization assay (Figure S3D, E), consistent with prior experiments placing the KH2F1–KREH2 interface in the N-terminal region of KREH2 (Kumar et al. 2019).

The second helicase cofactor, KH2F2, is homologous to fumarylacetoacetate hydrolase (FAH)-like domain-containing enzymes (AF-Q584U2-F1-v6, Figure 1A, D, S4A) (Hernandez et al. 2010; Kumar et al. 2016), which catalyze hydrolysis, decarboxylation, isomerization, or hydration reactions (Manjasetty et al. 2004; Tame et al. 2002). However, *T. brucei* KH2F2 lacks one of the three acidic residues required to coordinate a catalytic divalent metal ion and shows no homology in other active-site residues, suggesting compromised metal binding and therefore impaired catalytic activity (Figure 1E, F, S4A) (Izumi et al. 2007; Weiss et al. 2018, 2020). AlphaFold3 predicts that KH2F2 interacts with the dsRBD6 domain of KREH2, which we confirmed by a relocalization assay (Figure S4B, C); dsRBD6 is also known to bind RNA (Hernandez et al. 2010). The positively charged RNA-binding surface of dsRBD6, including the RNA-binding residue K1078 (Madina et al. 2015; Kumar et al. 2019), lies opposite the predicted KH2F2 interaction interface, suggesting that dsRBD6 could engage KH2F2 while simultaneously binding RNA. In conclusion, the AlphaFold models of KREH2 and its co-factors provide a basis for further detailed molecular analyses.

### KREH2 proximity proteomics links REH2C and RESC

To define the cellular context of the REH2C complex, we employed TurboID-based proximity labeling in living cells (Branon et al. 2018; Roux et al. 2012). We generated stable cell lines with C-terminal Turbo-ID tags on KREH2 or its co-factor KH2F1. A cell line expressing a mitochondrion-localized enhanced yellow fluorescent protein (MTS-eYFP) was used for data normalization. This engineering strategy generated mitochondrially localized expression of the fusion proteins (Figure S5A, B). The cellular proximity proteome revealed significant spatial association between KREH2 and its known cofactor KH2F1 (Figure 2A, Supplementary Table S7), validating our approach (Hernandez et al. 2010; Kumar et al. 2016, 2019). The proximity proteome of KH2F1 indicated a cellular context enriched for RESC subunits and general RNA processing factors (Figure 2B, Supplementary Table S8). The association with RESC is consistent with prior reports that suggested that this interaction may depend on RNA (Weng et al. 2008; Kumar et al. 2016).

**Figure 2.**
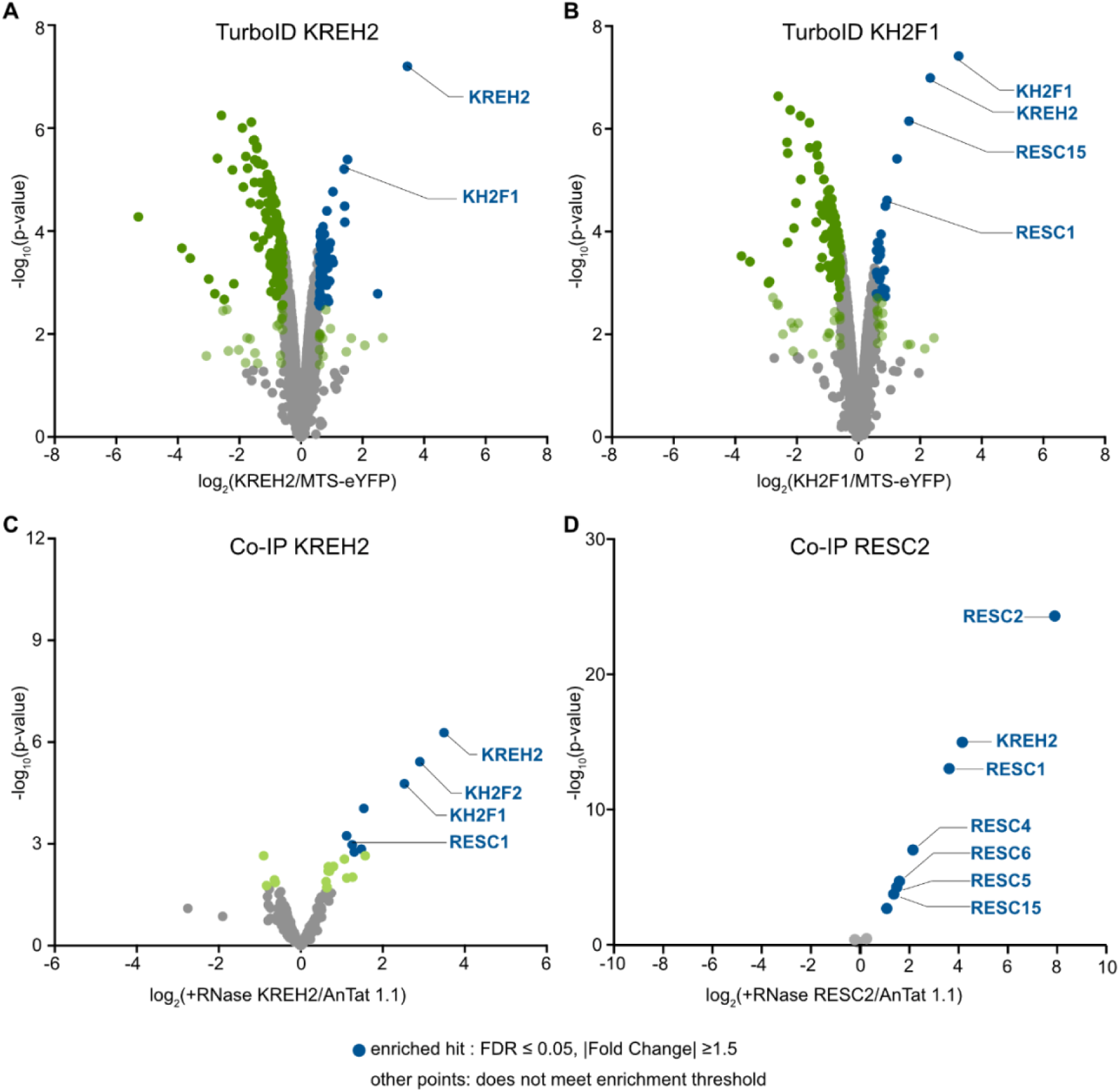
Proteins associated with KREH2, KH2F1, and RESC2 *in vivo*. **(A)** TurboID-tagged KREH2 compared to a MTS-eYFP control and (**B**) KH2F1 compared to MTS-eYFP. Volcano plots display log_2_(fold change) values calculated from DIA-NN intensities using a moderated t-test (limma package, (Ritchie et al. 2015). **(C)** Differential quantitative multiplexed mass spectrometry analysis of proteins co-purifying with endogenously tagged KREH2 compared to AnTat 1.1. (**D**) Differential quantitative multiplexed mass spectrometry analysis of proteins co-purifying with endogenously tagged RESC2 compared to AnTat 1.1. In C and D, samples have been treated with RNase. Volcano plots display log_2_(fold-change) values calculated from TMT intensities using a moderated t-test (limma package, (Ritchie et al. 2015). Significance threshold: Proteins were annotated as hits at a false discovery rate (FDR) ≤ 0.05 and |fold change| ≥ 1.5, and as candidates at FDR ≤ 0.2 and |fold change| ≥ 1.5. Independently, proteins were annotated as enriched if they were a hit or candidate with a positive fold change in the corresponding comparison of the tagged construct against untagged AnTat 1.1. Colours: dark blue, enriched hit; dark green, hit that is not enriched; light green, candidate that is not enriched; grey, all remaining proteins. Note that the y-axis shows −log_10_(p-value) whereas the hit/candidate cut-offs are applied to the FDR.

### KREH2 co-immunoprecipitates with RESC1-RESC2

To further refine the interactome of the KREH2 helicase and examine its binding partners, we performed co-immunoprecipitation experiments from cellular cultures with endogenously tagged proteins. Following a two-step affinity purification, including RNase treatment to disrupt RNA-dependent interactions, the co-immunoprecipitated proteins were identified by tandem mass tag (TMT) mass spectrometry. Consistent with the proximity proteome, co-immunoprecipitation detected a complex of KREH2 with KH2F1 and KH2F2, as reported by others, validating the experimental setup (Figure 2C, Supplementary Table S9) (Kumar et al. 2016). Bands corresponding to the expected sizes of the trimeric REH2C complex were detected by silver-stained PAGE analysis of the sample (Figure S6). The KREH2 co-immunoprecipitation interactome also indicated an interaction with RESC1, consistent with a prior study (Kumar et al. 2016). Notably, the RNase resistance suggests that this interaction may be direct. The co-immunoprecipitation interactome of RESC2, the obligatory binding partner of RESC1, which we tagged for cross-correlation, in turn showed enrichment of KREH2 and also additional RESC subunits (Figure 2D, Supplementary Table S10). Thus, the ensemble of proximity-labeling and co-immunoprecipitation experiments confirms the reported KREH2-KH2F1-KH2F2 complex and its association with RESC *in vivo*. Additionally, our data suggest RNase-resistant interactions between REH2C and RESC complexes, particularly with the RESC1-RESC2 heterodimer.

### An internal peptide of KREH2 directly binds to the RESC1 surface

The potential of direct, RNA-independent interactions between the REH2C complex and the RESC1-RESC2 heterodimer motivated an AlphaFold-multimer analysis (Jumper et al. 2021) and evaluation of the interaction using isothermal titration calorimetry (ITC). The binding experiment indicated a high-affinity equimolar interaction (Kd=220±20 nM; N=0.979±0.006) between the conserved KREH2 peptide 560–579 and RESC1-RESC2 (Figure 3A, B). Next, we determined the cryo-EM structure of this complex, achieving an overall resolution of 2.9 Å and a local resolution of 3.0–3.5 Å for the KREH2 peptide (Figure 3C, D, S7, Table 1). The structure of the RESC1-RESC2 complex is similar to previously reported cryo-EM structures (RMSD: 0.398 Å for RESC1 and 0.390 Å for RESC2 to PDB:8CDP) (Dolce et al. 2023; Liu et al. 2023); however, the side-chain definition has improved owing to the higher resolution. In brief, the RESC1-RESC2 heterodimer adopts a dumbbell-like shape, with each subunit forming one lobe. Each subunit adopts an imperfect β-barrel flanked by α-helices, with dimerization mediated by a domain-swapped β-hairpin. Map density accommodates KREH2 residues 565–571 (RRVAKRA) with the position closely resembling the AlphaFold3 prediction (peptide main chain RMSD: 1.790 Å for 28 atom pairs) (Figure 3E, S8A, B) (Abramson et al. 2024). The basic peptide binds to the exterior surface of the RESC1 barrel near β-strands β14 and β15, which form a negatively charged binding surface (Figure 3F). The interaction is primarily electrostatic, with several potential hydrogen bonds further stabilizing the complex (Figures 3G, S8C–E), rationalizing the high binding affinity despite the limited contact area. Inverse-charge substitution of interface residues reduced affinity (R565E and K569E) or abolished binding (R566E and R570E), respectively, demonstrating that these residues are essential for complex formation (Figure 3A, B). To ensure complete disruption of the full predicted basic contact patch (residues 564–570), a quadruple-mutant variant incorporating flanking residue K564 (K564E/R566E/K569E/R570E) was also evaluated and confirmed to abolish binding (Figure 3A, B). In conclusion, affinity measurements and cryo-EM analyses demonstrate that the examined KREH2 peptide directly binds the RESC1-RESC2 complex with high affinity via conserved electrostatic interactions.

**Figure 3.**
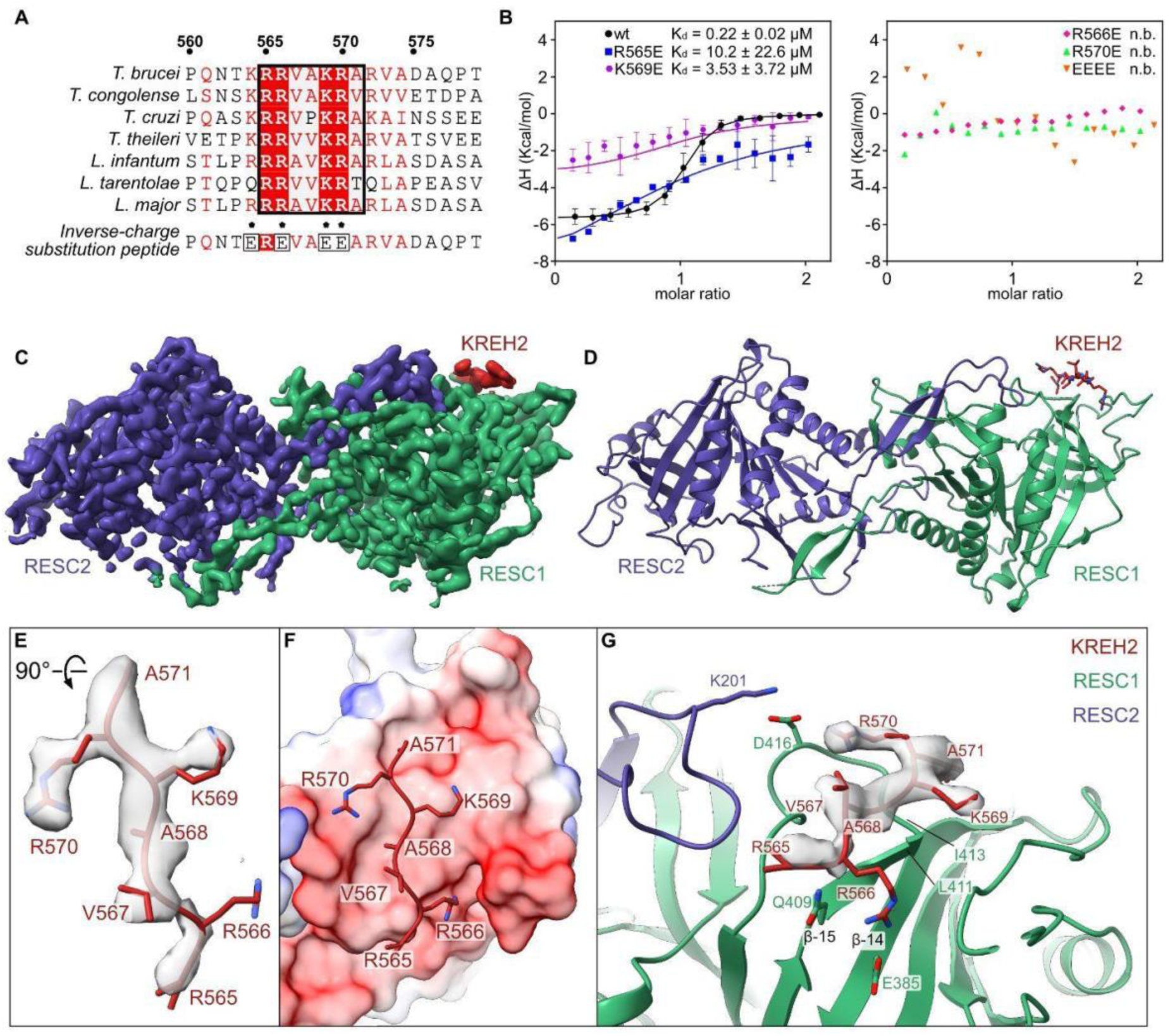
The direct protein-protein interaction of a KREH2 peptide with RESC1-RESC2. (**A**) Multi-species sequence alignment of the KREH2 binding peptide. Residues resolved in the cryo-EM map are highlighted in a black box. 100% conserved residues are shown in red boxes, and 70% conserved residues are shown in red. Residues substituted with amino acids of inverse charge for experimental analysis are marked with black stars. The quadruple-mutant peptide is also shown. (**B**) ITC binding curve of RESC1-RESC2 with KREH2 peptides. The plot shows the change in heat (kcal/mol) as a function of molar ratio. The data represent the mean of three independent experiments (n = 3) fitted to a single-site binding model for wt. Two independent experiments (n =2) were performed for mutant peptides with reduced binding, and a single experiment was performed for the mutant peptides with no binding. n.b.= no binding. (**C**) Cryo-EM map of the RESC1-RESC2 heterodimer (green and purple, respectively) bound to the KREH2 peptide (red). (**D**) Cartoon model of RESC1-RESC2-KREH2; colors as in C. (**E**) Cryo-EM density of the KREH2 peptide with cartoon and stick model. (**F**) The interaction interface between RESC1 and the KREH2 peptide. RESC1 as surface coloured by coulombic electrostatic potential (blue positive and red negative) calculated by ChimeraX; KREH2 as a cartoon and stick model. (**G**) Zoomed in view of the interaction interface between RESC1 and KREH2 peptides. The DeepEMhancer postprocessed cryo-EM map is shown at 0.12 level.

**Table 1:**
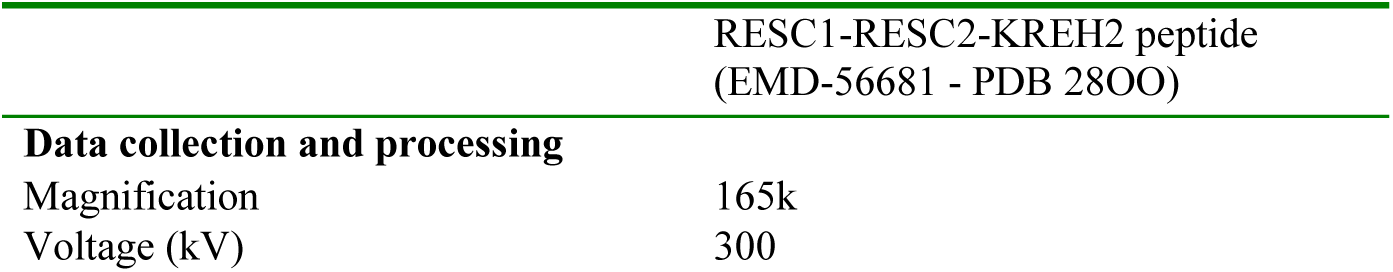
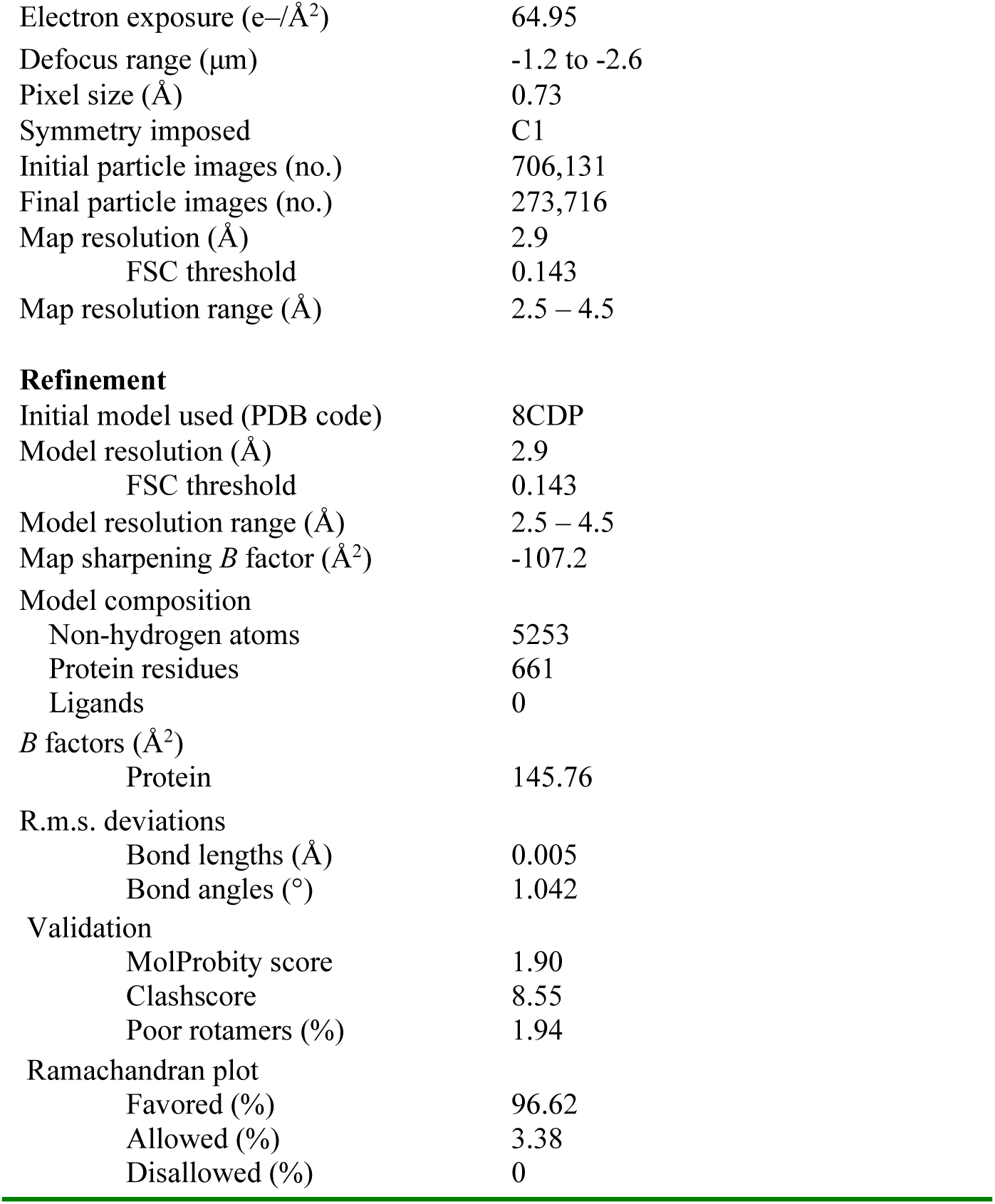
Cryo-EM data collection, refinement and validation statistics.

### Disruption of the KREH2-RESC1 interface does not lead to growth and RNA editing defects *in vivo*

To assess the functional importance of the KRRVAKR motif (positions 564-570) in KREH2, we established a genetic complementation system in bloodstream-form (BSF) *T. brucei*. We generated a single-allele knockout (SKO) of KREH2 and introduced a tetracycline (tet)-inducible RNA interference (RNAi) construct targeting the endogenous 3′ untranslated region (UTR) of the remaining allele. The parental SKO/RNAi clonal cell line proliferated at normal rates in the absence of induction. Upon tet induction, however, KREH2 protein levels declined by 80% by day 1, and difference in parasite growth emerged by day 2 (Figure 4B, S9A). To complement this growth defect, we expressed RNAi-resistant ectopic KREH2 variants - either wild-type or the inverse-charge tetra-substitution mutant K564E/R566E/K569E/R570E - from the tubulin locus in the parental cell line. Following tet-induction, exclusive expression of wild-type KREH2 fully restored normal growth (Figure S9B). Independent clonal lines of the K564E/R566E/K569E/R570E mutant also revealed that exclusive expression of the mutant KREH2 could complement cell growth (Figure 4B, S9C-F). All subsequent studies of editing were conducted 24 hours post-induction when secondary effects of KREH2 knockdown on the abundances of other editing complex components, e.g., KREPA1 protein (RECC complex), were not observed (Figure S9G,H). Western blot analyses showed comparable steady-state levels of KREH2 wild-type and mutant variants, suggesting that the sequence mutagenesis had no visible effect on protein stability (Figure 4A). To further examine any possible functional effects of the K564E/R566E/K569E/R570E mutant on RNA editing, we performed RT-qPCR on nine of the 12 mitochondrial editing targets both in wild-type and mutant variants in our complementation system (Figure 4C). These editing panels showed that the K564E/R566E/K569E/R570E substitutions in KREH2 did not result in gross steady-state editing defects and allowed the cells to generate fully edited messages. We cannot rule out the possibility of more subtle editing defects that are not detectable by this qPCR assay. However, this data indicates that the observed KREH2 KRRVAKR motif-RESC1 interaction is not absolutely essential for editing function, and that perhaps additional complementary interactions between KREH2 and/or its co-factors, and RESC1-RESC2 are able to maintain function.

**Figure 4:**
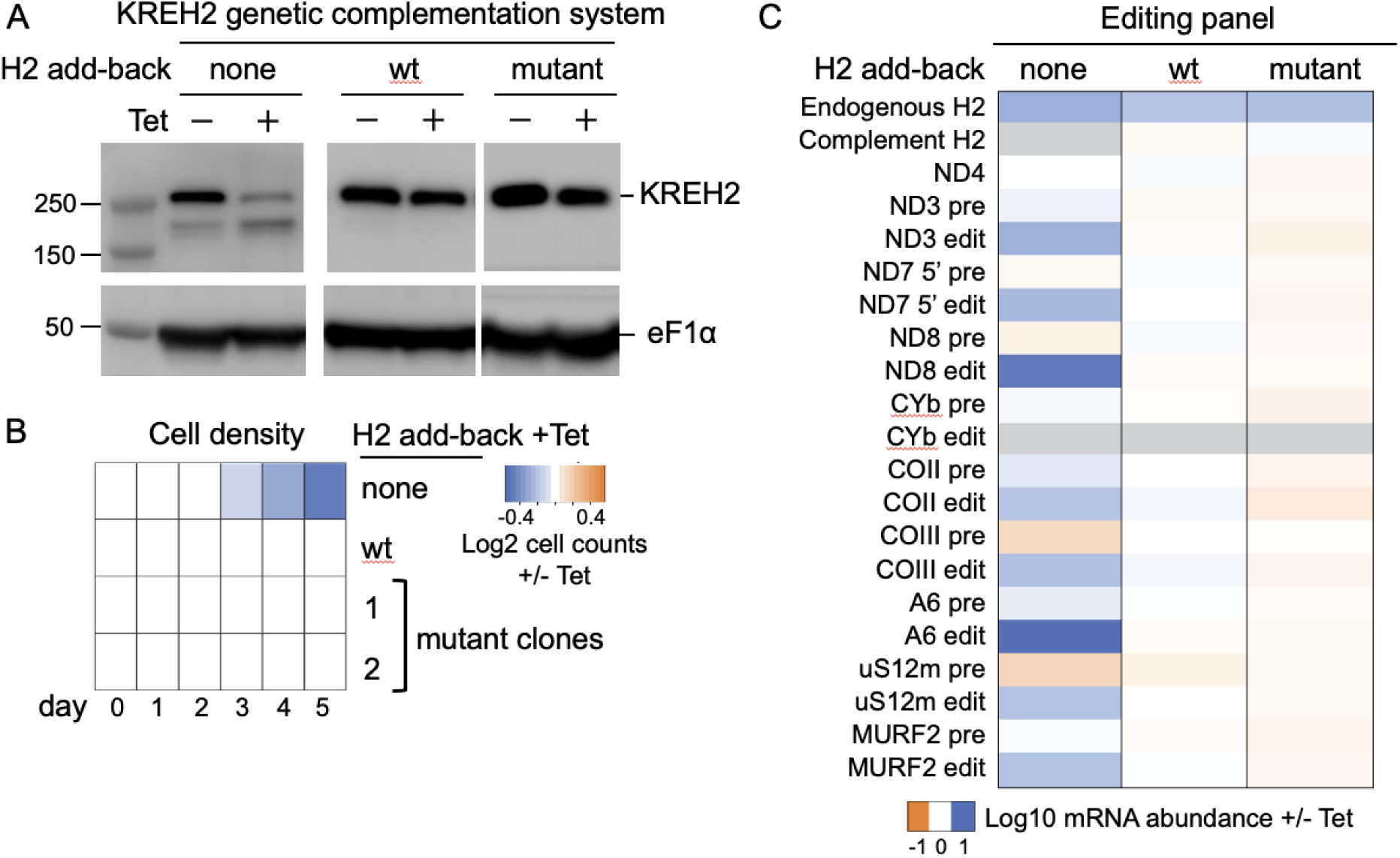
KREH2 KRRVAKR motif-RESC1 interaction is not essential for editing function. (**A**) KREH2 complementation system: BSF KREH2 SKO/RNAi cell lines that lack (none) or contain a constitutively expressed wt or K564E/R566E/K569E/R570E mutant allele were grown in -tet versus +tet to induce endogenous KREH2 knockdown. Protein lysates were analyzed via western blot using antibodies against KREH2 and eF1a as a loading control. (**B**) Cell lines in panel A were grown in +tet to induce endogenous KREH2 knockdown. Heat map displays the log2 ratio of cumulative growth every day for five days. Reduction or increase in growth upon knockdown, or exclusive expression of WT or mutated alleles are indicated by blue or orange color scale, respectively. Data are representative of a minimum of four independent experiments. (**C**) Consequences to mRNA editing upon KREH2 loss and complementation by constitutively expressed wt or K564E/R566E/K569E/R570E mutant alleles in BSF cells as shown by quantitative RT-PCR analyses. Relative endogenous KREH2, complementary wt or mutant KREH2, never-edited ND4, and pre- and fully-edited mRNA abundances in cells grown for 24 hours in the presence of tet (endogenous KREH2 knockdown via RNAi and exclusive expression of wt or K564E/R566E/K569E/R570E mutant) vs. the absence of tet (endogenous KREH2 expressed) were determined by qRT-PCR. For each target amplicon, the relative change in RNA abundance was determined using the mRNA encoding telomerase reverse transcriptase (TERT) as an internal control. Data are shown as means from at least two independent experiments (means ± SEM are shown in Figure S9I). Grey boxes indicate that the mRNAs were not detected in cells grown in both minus tet and plus tet.

## Discussion

RNA helicases are dynamic choreographers of ribonucleoprotein remodeling across diverse cellular pathways, as disparate as innate immune signaling, ribosome biogenesis, and pre-mRNA splicing (Jarmoskaite and Russell 2014). Growing understanding of kinetoplastid RNA editing and its associated complexes indicates that the editing cycle entails extensive de novo assembly and remodeling of large, modular assemblies, with several RNA helicases participating as facilitators. Here, we identify a previously unrecognized direct interaction of the mitochondrial RNA editing helicase KREH2 with the obligate RESC1–RESC2 heterodimer anchored at the 5′ ends of all gRNAs, suggesting an important functional connection between KREH2 and gRNAs (Figure 5).

**Figure 5:**
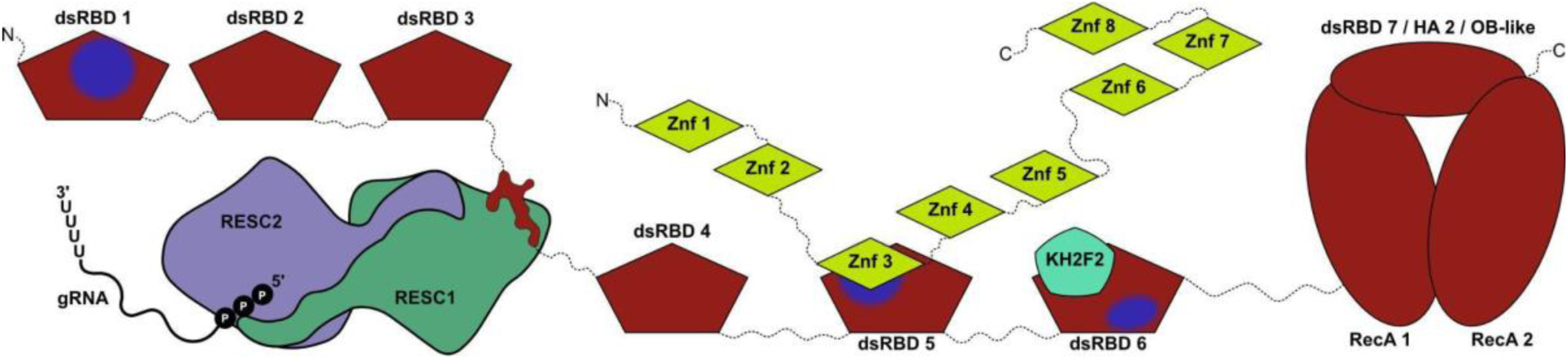
Model of KREH2 bound to gRNAs via RESC1-RESC2 together with KH2F1 and KH2F2. Based on AlphaFold predictions and available experimental data. KREH2 in red, KH2F1 in light green, KH2F2 in light blue, RESC1 in dark green, RESC2 in dark purple, and gRNA in black. Positively charged surfaces of the dsRBD domains of KREH2 are highlighted in blue.

The RESC1-RESC2 module is central to RNA editing, recognizing the 5′-triphosphate moiety of gRNAs through interaction within a positively charged tunnel in RESC2 (Dolce et al. 2023; Liu et al. 2023; Weng et al. 2008), thereby protecting the gRNA pool from degradation (Dolce et al. 2023; Liu et al. 2023). Other functions of RESC1 were unknown, but it was speculated that it might provide additional surfaces as platforms for recruiting complexes that interact with gRNAs (Dolce et al. 2023). Here, we identify KREH2 as a direct interactor of RESC1 using cellular proximity labeling and purification of the complex from RNA-treated cell lysate combined with mass spectrometry-based identification of interaction partners. We provide evidence of the direct binding through structural biochemistry with a recombinant protein. In addition to its ability to autonomously bind gRNAs, the RESC1–RESC2 heterodimer was reported to participate in two mutually exclusive gRNA-bound RESC variants: canonical RESC, comprising obligate RESC1-RESC2, and RESC5 and RESC6 subunits, other RESC proteins, and fully-edited mRNAs; and the alternative RESC*, which lacks RESC5–RESC6 and minimally contains RESC1–RESC2. Both canonical RESC, an editing competent variant, and the minimal RESC*, were shown to transiently and stably associate with REH2C, respectively (reviewed in Cruz-Reyes et al. 2018; McDermott et al. 2026; Kumar et al. 2016). Recent cryo-EM particle reconstructions also revealed a variety of heterogeneous RESC assemblies: ‘RESC-A’, which exclusively binds gRNA and potentially serves to sequester gRNAs for their storage, while ‘RESC-B’ and ‘RESC-C’ both bind pre-mRNA and gRNA (Liu et al. 2023). RESC-A contains RESC5–RESC6 and RESC1–RESC2, while RESC-B and RESC-C both carry RESC5–RESC6 but lack RESC1–RESC2 (Liu et al. 2023). Within RESC-A, the RESC1-RESC2 dimer has been reported to engage the 5′-triphosphate of the gRNA (Dolce et al. 2023; Liu et al. 2023), RESC3 and RESC4 appear to serve scaffolding roles, and RESC5 and RESC6 bind the gRNA poly(U) tail (Liu et al. 2023). In contrast, RESC-B contains RESC5 to RESC14 proteins, with the exception of RESC12/12A, with RESC5-RESC6 remaining bound to the gRNA poly(U) tail (Liu et al. 2023). RESC-A is thought to be a gRNA-stabilizing complex, while RESC-B is suggested as a probable substrate for the catalytic RECC (Liu et al. 2023), scaffolding mRNAs and gRNAs during editing. Both our proteomics approaches, in combination with prior studies, support the existence of a minimal KREH2-RESC1-RESC2 complex that could possibly overlap with the previously described RESC* assembly (McDermott et al. 2026; Kumar et al. 2016), but that is independent of RESC-A or RESC-B contexts, or the editing-competent RESC (Liu et al. 2023; Ammerman et al. 2012; Kumar et al. 2016). The superposition of our complex presented here with the RESC-A cryo-EM structure indicates that KREH2 peptide binding is, in principle, sterically compatible with the additional RESC subunits present in the known larger complexes (Figure S8F). Thus, the absence of other RESC components in our proteomics datasets cannot be readily explained by the available structural information and requires further investigation.

KREH2, a DEAH/RHA-family helicase, shares homology with well-characterized RNA helicases such as MLE, Prp2, Prp43, and DHX15; these remodel RNPs in gene silencing and pre-mRNA splicing. In MLE, the two N-terminal dsRBDs function in a context-dependent manner, acting both cooperatively and independently to bind RNA and regulate subcellular localization (Prabu et al. 2015; Jagtap et al. 2023; Maenner et al. 2013). This homology suggests that KREH2 may similarly remodel RNA or RNA-protein complexes across different contexts. Uridine insertions dominate kinetoplastid RNA editing, rendering pre-edited mRNAs G-rich and prone to forming G-quadruplexes (Leeder et al. 2016). Given that DEAH/RHA-family helicases such as DHX36 and DHX9 can resolve G4 structures (Tippana et al. 2019; Chakraborty and Grosse 2011), KREH2 may similarly promote efficient editing by remodeling these structures, a hypothesis that remains to be tested. The kinetoplast RNA-editing helicases KREH1 and KREH2 perform distinct roles. Unlike KREH2, KREH1 locally unwinds dsRNA, including mRNA:gRNA duplexes and gRNA secondary structures, to promote editing initiation in pan-edited transcripts (Dubey et al. 2023). KREH1 association with the editing machinery is RNA-dependent and is mediated by its C-terminus (Yelmar et al. 2026). Furthermore, another DEAH-box helicase, KREH3, has been identified in KREH2 purifications (Hernandez et al. 2010), and the ABC ATPase KRBP72 can interact with A6 pre-mRNA (Dubey et al. 2025); further research will clarify the distinct roles of these potential RNA remodeling factors.

The proteomics and structural analyses presented here corroborate the existence of a stable REH2C complex consisting of KREH2, KH2F1, and KH2F2 (Kumar et al. 2019, 2016; Madina et al. 2015; Hernandez et al. 2010; McDermott et al. 2026). Previously reported RNA-dependent interactions with RECC and RESC assemblies, along with association with mRNAs across multiple editing stages (Madina et al. 2015; Meehan et al. 2023; Kumar et al. 2016; Meehan et al. 2024), suggest that the direct KREH2–RESC1 interaction represents only one facet of a broader holo-editosome interaction landscape of the helicase, consistent with the sophisticated multidomain architecture of KREH2. Additional molecular contacts could be mediated by the multiple dsRBDs (Kumar et al. 2019; Gleghorn and Maquat 2014; Hitti et al. 2004; McDermott et al. 2026), by the disordered regions of KREH2, or by the co-factors KH2F1 and KH2F2. In particular, KH2F1, whose zinc finger domains are predicted to bind RNA, likely contributes to RNA-mediated complex formation. RNAi-mediated KH2F1 depletion impaired RNA editing (Kumar et al. 2016); its role could be reminiscent to G-patch proteins, which regulate DEAH/RHA-family helicases in the spliceosome to enhance helicase activity and substrate specificity (Studer et al. 2020; Cruz-Reyes et al. 2016; Tanaka et al. 2007; Christian et al. 2014). However, further molecular characterization of KH2F1 and KH2F2, along with structural elucidation of the full REH2C complex, will be required to clarify their roles.

To conclude, this study advances our mechanistic understanding of kinetoplastid RNA editing by demonstrating that KREH2 engages guide RNAs via the RESC1–RESC2 complex. The findings highlight the dynamic RNA remodeling in RNA editing through helicases, resembling aspects of pre-mRNA splicing.

### Limitations of the study

For the TMT-MS interactome study, a non-clonal cell population was used. Additionally, the interactome analysis was performed using technical triplicates derived from the same cell lysate; therefore, the reported FDRs reflect technical rather than biological reproducibility. In the TurboID experiment, C-terminal tagging of KREH2 likely limited detection of proteins near its N-terminus, given its large, flexible, multidomain structure; however, N-terminal tagging perturbed mitochondrial localization, according to the TrypTag database (Billington et al. 2023), and was therefore not considered. TurboID labeling may miss proximal proteins that lack accessible lysine residues for covalent attachment, potentially due to steric hindrance within multiprotein assemblies. Alternatively, low expression levels or tag-induced alterations in the folding of the bait fusion protein may result in reduced detection of proximal proteins. Furthermore, small proteins may escape detection by mass spectrometry. We reconstituted a minimal RESC1-RESC2-KREH2 peptide complex; how the full-length helicase would align in three-dimensional space remains unclear.

## Data Availability

The cryo-EM map is available at EMDB: EMD-56681 and the model is deposited at PDB: 28OO/pdb_000028oo. The mass spectrometry proteomics data have been deposited to the ProteomeXchange Consortium via the PRIDE partner repository with the dataset identifier PXD081786.

## Acknowledgements

We acknowledge Martin Pelosse for support in using the Eukaryotic Expression Facility at EMBL Grenoble, and Romain Linares and Iskander Khusainov for support in using the EM Facility at EMBL Grenoble. We thank the European Synchrotron Radiation Facility for providing beam time on CM02 and thank Félix Weis for his assistance with data collection; the data from CM02 are in the ESRF data portal doi.org/10.15151/ESRF-ES-2265597421. This work used the platforms of the Grenoble Instruct-ERIC center (ISBG; UAR 3518 CNRS-CEA-UGA-EMBL) within the Grenoble Partnership for Structural Biology (PSB), supported by FRISBI (ANR-10-INBS-0005-02) and GRAL, financed within the University Grenoble Alpes graduate school (Écoles Universitaires de Recherche) CBH-EUR-GS (ANR-17-EURE-0003). We are grateful to Caroline Mas for assistance with biophysical experiments, especially ITC. We thank EMBL Heidelberg, Jennifer Schwarz, and Frank Stein from the EMBL Proteomics Core Facility for their support with the mass spectrometry data acquisition and analysis. The project was previously supported by a grant from the French Agence Nationale de la Recherche to E.K. (ANR-20-CE11-0016). The lab of Eva Kowalinski is supported by an ERC grant, TRANSPLIC, 101170068, funded by the European Union. Views and opinions expressed are, however, those of the authors only and do not necessarily reflect those of the European Union. Neither the European Union nor the granting authority can be held responsible for them. The Cruz-Reyes lab is supported by an NIH R01 AI184920 to JCR. The authors thank Jerusha Abraham and Christopher Lo at Texas A&M AgriLife for assistance with transfection, generation of clonal cell lines, and growth curves. The lab of Suzanne McDermott is supported by NIH R01 AI184920 and Seattle Children’s Research Institute seed funds. The authors thank their lab members for active discussions and constructive comments throughout the project.

## Author contributions

EK study lead, design and funding acquisition. JCR funding acquisition, figure design. RY and LD carried out the AlphaFold analysis. LT and JC carried out ITC experiments. RY carried out the co-immunoprecipitation experiment. BPG carried out the Turbo-ID experiment. RY and LT prepared the cryo-EM sample. LD and EK processed and interpreted the cryo-EM data. LOH generated the mutant KREH2 plasmid construct, generated one *T. brucei* clonal cell line of mutant KREH2 add-back, carried out WT vs. mutant KREH2 genotyping, growth analyses, RNA extraction, cDNA synthesis for BioMark, and western blot analyses from parental KREH2 SKO/RNAi, WT, and mutant KREH2 add-back *T. brucei* cell lines. ZG generated the KREH2 SKO plasmid construct. CG generated the KREH2 SKO *T. brucei* cell line, one *T. brucei* clonal cell line of mutant KREH2 add-back, genotyping, growth analyses, and generation of figures. SKS generated the WT KREH2 plasmid construct and the *T. brucei* WT KREH2 add-back cell line, transfected the KREH2 RNAi construct into *T. brucei* KREH2 SKO cells, growth analyses, genotyping, and western blot analyses. TR carried out RNA extraction from WT vs. mutant KREH2 T. brucei cell lines. SMM carried out WT vs. mutant KREH2 *T. brucei* cell line validation, growth analyses, and BioMark qRT-PCR analyses of mitochondrial transcripts. RY, LD, and EK wrote the manuscript with the contribution of CG, JCR, and SMM.

