## Supplementary Material for "The RNA helicase KREH2 directly binds the RESC1-RESC2 complex in trypanosomatid mitochondrial RNA editing"

### Supplementary Methods

#### Streptavidin blotting and imaging

To detect biotinylation on a fluorescence microscope in fixed cells, cells expressing KREH2::TurboID-HA, KH2F1::TurboID-HA, and MTS-TurboID-HA-eYFP were chemically fixed, permeabilized, and stained with streptavidin-conjugated AlexaFluor 594 and DAPI. Streptavidin imaging was performed as previously described in (Odenwald et al. 2024), except that streptavidin labeled with AlexaFluor® 594 (ThermoFisher) was used. Fluorescence images were acquired on a confocal microscope LSM900 (Zeiss) equipped with a Plan-Apochromat 63×/1.40 NA oil immersion objective, using the 405 and 561 nm laser excitation for DAPI and AlexaFluor® 594, respectively. Fiji was used for figure generation (Odenwald et al. 2024; Schneider et al. 2012).

To detect biotinylation in a western blot, parental cells and cells expressing either KREH2::TurboID-HA, KH2F1::TurboID-HA and MTS-TurboID-HA-eYFP were harvested by centrifugation (1,400 g, 10 min, 27 °C) and washed once in serum-free SDM79 medium (1,400 g, 5 min, 27 °C) before resuspension in a 2× SDS loading buffer for a concentration of $5x10^5$ cells/µl and incubation at 95 °C for 5 min. Lysates were resolved on 4-20% gradient SDS–PAGE gels ($10^6$ cells/lane) and proteins transferred onto a PVDF membrane in Tris-glycine-ethanol transfer buffer using a wet-transfer system. The membrane was blocked in 5% (w/v) milk in PBS for one hour and subsequently incubated with IRDye® 680RD Streptavidin (LICORbio; 1:1000) and anti-HA (Cell Signaling, C29F4; 1:5000) for 16 h, followed by incubation with IRDye® 800CW Goat Anti-Rabbit IgG (LICORbio; 1:20,000) for one hour. Streptavidin and antibodies were diluted in PBS-T (0.1% Tween-20 (v/v) in PBS). Washing steps were performed 4 times, each for 5 minutes in an excess of PBS-T after each incubation. The last wash of the secondary antibody was carried out in PBS. Membranes were dried between two filter papers and imaged on an Odyssey® M Imaging System (LICORbio) using the 700 and 800 nm channels.

#### Relocalization (ReLo) assay

To experimentally assess the AlphaFold3-predicted interactions, we used a relocalization assay (Salgania et al. 2024) adapted for HEK-293T cells. Briefly, HEK-293T cell lines constitutively expressing an anti-mCherry nanobody fused to pleckstrin homology (PH) domain were established. These were co-transfected with two psLIB vectors (Pelosse and Marcia 2025), one transiently expressing a mCherry-fused protein, anchored to the inner plasma membrane via the nanobody, and the other GFP-fused potential interacting protein, whose re-localization to the PM is indicative of interaction. We tethered mCherry-tagged constructs of the KH2F1 Znf2 (residues 101–165) and Znf3 (residues 223–265), or the Fumarylacetoacetase-like domain of KH2F2 (residues 11–267) to the plasma membrane, and assessed the re-localization of GFP-tagged constructs flanking KREH2 dsRBD5 (residues 816–959) and dsRBD6 (residues 967–1116). Transfections and cell line generation were carried out as described in (Pelosse and Marcia 2025).

Fluorescence images were acquired on a confocal microscope LSM900 (Zeiss) equipped with a Plan-Apochromat 63×/1.40 NA oil immersion objective, using 488 and 561 nm laser excitation for GFP and mCherry, respectively. Fiji (Odenwald et al. 2024; Schneider et al. 2012) was used for figure generation and to generate line-intensity profiles. Lines were manually drawn on merged images, and intensity values were extracted from each individual channel. For each image and channel, intensities were normalized to the maximum intensity measured within the same profile. To account for differences in line length between images, profiles were resampled by linear interpolation to 101 equally spaced positions spanning 0-100% of the line length.

### Supplementary Figures

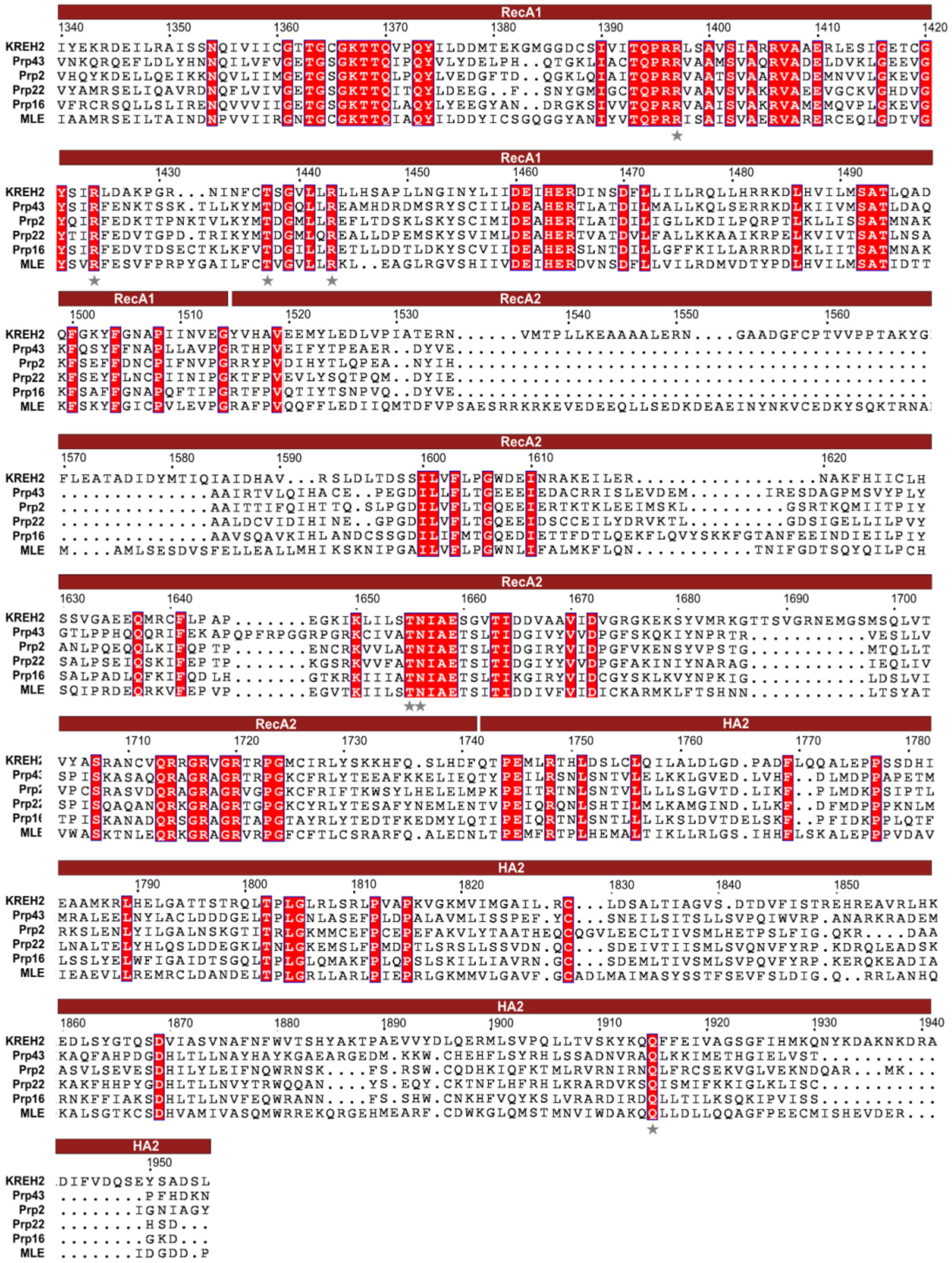


**Figure S1: Multiple sequence alignment of the KREH2 C-terminal domains.** Clustal Omega alignment of *T. brucei* KREH2 C-terminal domains, RecA1, RecA2 and Helicase-associated domain 2 (HA2) with other DExH box helicases, Prp43 (*S. cerevisiae*; P53131), Prp2 (*S. cerevisiae*; P20095), Prp22 (*S. cerevisiae*; P24384), Prp16 (*S. cerevisiae*; P15938), MLE (*D. melanogaster*; P24785), visualized with ESPript 3.2. 100% conserved residues are shown in white characters on a red background. Residues in Prp43 previously reported to interact with RNA in the RNA-binding tunnel and hook-turn region, and conserved across DExH-box helicases, including KREH2, are indicated by grey stars beneath the alignment.

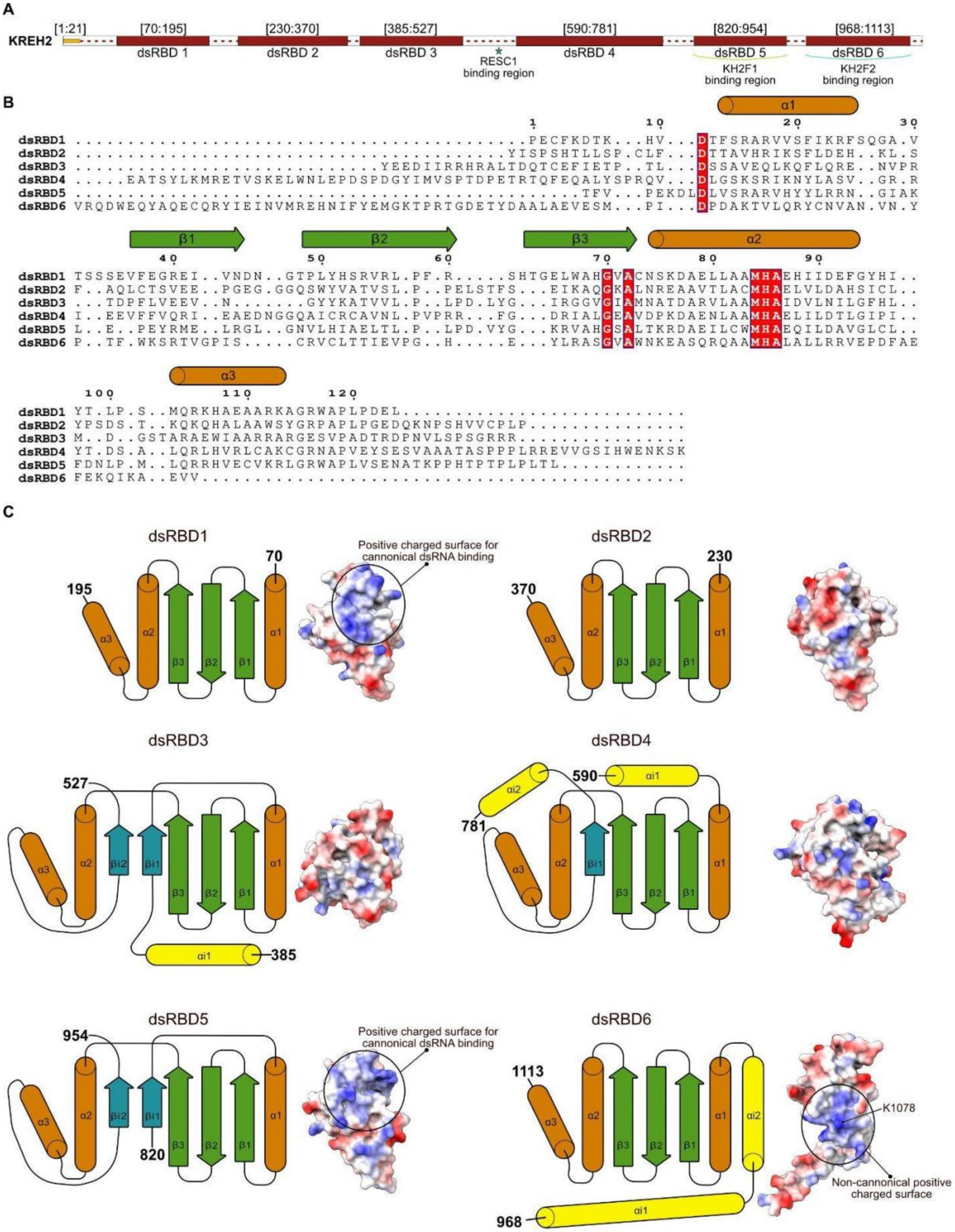


**Figure S2: The KREH2 N-terminal dsRBDs.** (**A**) Domain organization of the N-terminal region of KREH2, based on the AlphaFold predictions. dsRBDs are shown as boxes, the mitochondrial targeting sequence in yellow, and the disordered regions as dashed lines. The predicted interfaces towards the helicase co-factors are annotated. (**B**) Multiple sequence alignment based on the structures of the 6 dsRBDs generated with the PDBeFold server and visualized with ESPript 3.2. Strictly conserved residues (100% identity) are shown as white characters on a red background. Secondary-structure elements derived from the AlphaFold prediction of dsRBD1 above the alignment. (**C**) Topology diagrams of the KREH2 dsRBDs, with the canonical helices and sheets colored in orange and green, and the insertions colored in yellow and slate. Surface models with electrostatic surface potential (red negative; blue positive) of the dsRBDs are shown. Positively charged surfaces that potentially can bind nucleotides are annotated.

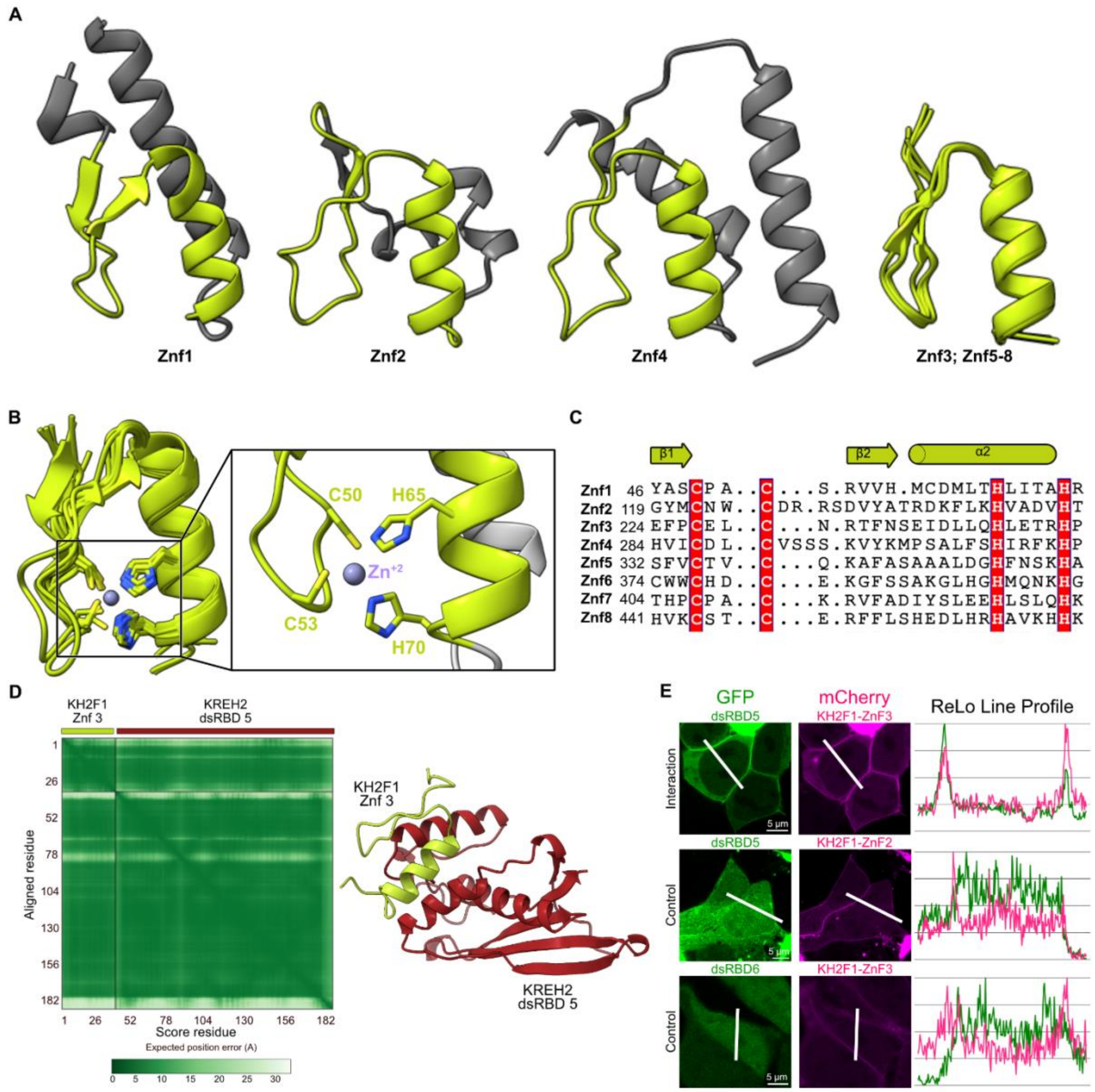


**Figure S3: The KREH2 co-factor KH2F1.** (**A**) AlphaFold predicted structure of KH2F1 Zinc fingers (Znf) as cartoon representations. The common Zn2+-coordinating β-β-α structure is shown in light green, and additional helices in grey. The four representative Znf architectures shown are grouped according to structural features of the Znf core. Znf1, Znf2, and Znf4 contain additional structural features beyond the canonical C2H2 core, whereas Znf3 and Znf5–8 share the canonical C2H2 core architecture. This structural grouping is distinct from the three ZnF groups described in the Results, which were defined based on the PAE plots. (**B**) Structural superposition of Zn2+ coordinating core structure of all eight KH2F1 Znfs in cartoon representation. Side chains of Zn2+-coordinating residues are shown as sticks in heteroatom coloring. Zoomed view of Zn2+-coordinating residues from Znf1, highlighting conserved spatial arrangement. Zn2+ ions were modeled based on structural alignment using the Znf5 domain of Xenopus laevis transcription factor IIIA (PDB: 1TF6) as a template. (**C**) Structural multiple sequence alignment of all Znfs generated with mTM-align and visualized with ESPript 3.2. The 100% conserved Zn2+ coordinating residues are shown as white characters on red background. Secondary-structure elements derived from the AlphaFold-predicted Znf1 structure are shown above the alignment (α-helices as cylinders and β-strands as arrows), with residue numbering corresponding to the start of each Zn finger. (**D**) AlphaFold3 prediction of the interaction interface between Znf3 of KH2F1 and dsRBD5 of KREH2, with Predicted Alignment Error (PAE) plot and predicted model presented in cartoon. (**E**) Relocalization (ReLo) assay in mammalian cells probing the interaction between KH2F1 Znf domains and KREH2 dsRBDs. (Left and Center) Confocal microscopy images of cells co-transfected with a cytoplasmic 'prey' domain (GFP-dsRBDs) and a plasma-membrane-targeted 'bait' domain (mCherry-Znf). We detect interaction by recruiting GFP-labeled prey to the plasma membrane, as observed for dsRBD5 with KH2F1-Znf3 (top row). Non-interacting pairs (bottom two rows) show cytoplasmic localization of the GFP-labeled prey. A white line on each image marks the path used for line profile analysis. Scale bars are 5 µm. (Right) ReLo line profiles showing the normalized fluorescence intensity of the GFP (green) and mCherry (magenta) signals along the indicated lines. Co-localization of both peaks indicates an interaction.

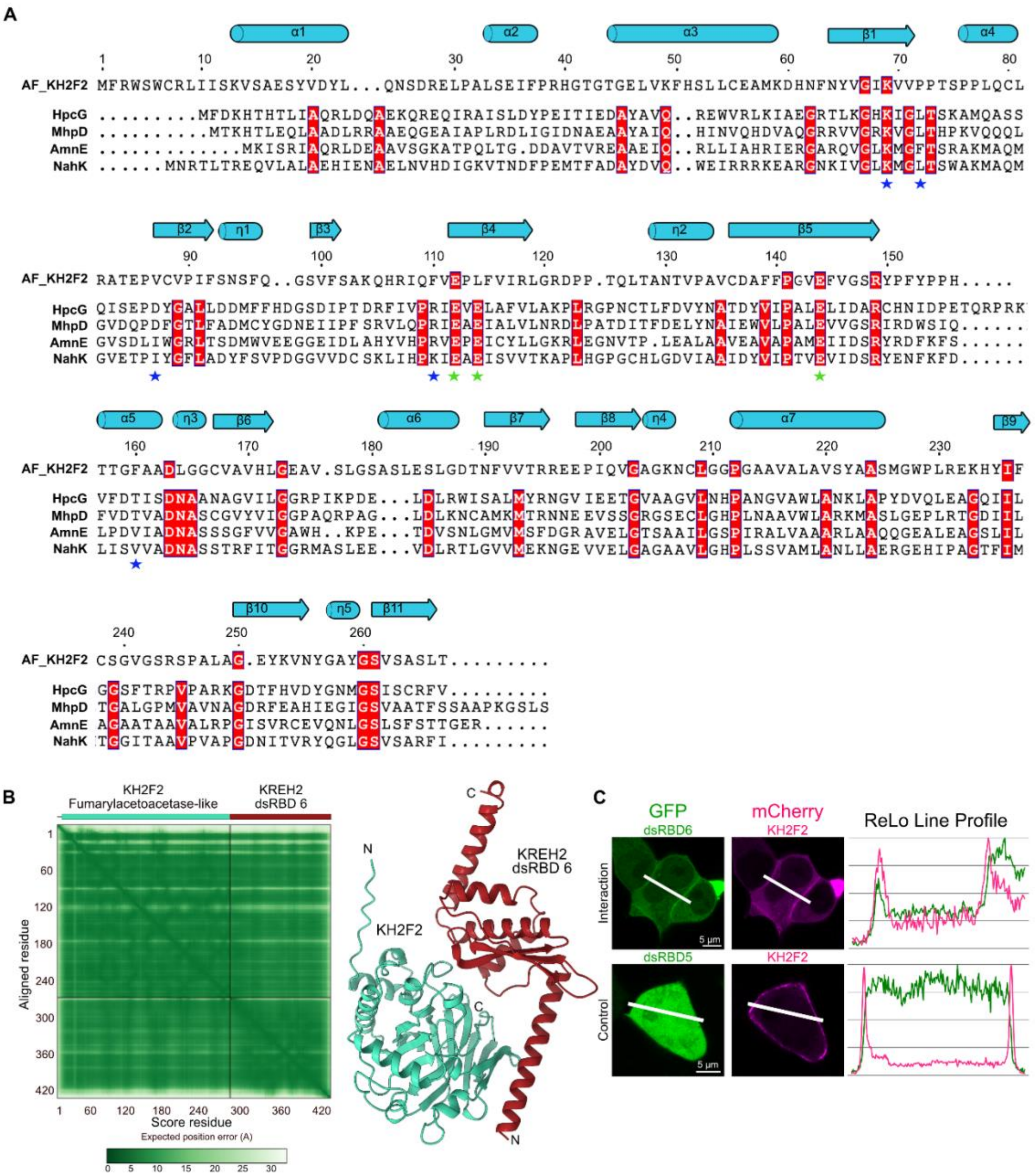


**Figure S4: The KREH2 co-factor KH2F2.** (**A**) Multiple sequence alignment of T. brucei KH2F2 and FAH-domain containing homologs, HpcG (2EB5), AmnE (8IH6), NahK (5D2F), and MhpD (1SV6) from *E. coli* and Pseudomonas spp. was generated with ESPript 3.2. Strictly conserved residues (100% identity) as white characters on red background. Three acidic divalent metal ion-coordinating residues conserved across the enzymatic homologs are highlighted with a green star; residues contributing to the active site are indicated with blue star. Secondary-structure elements derived from the AlphaFold model of KH2F2 are shown above the alignment (α-helices as cylinders and β-strands as arrows), with residue numbering corresponding to the KH2F2 sequence. (**B**) AlphaFold3 prediction of the interaction interface between KH2F2 and dsRBD6 of KREH2, with PAE plot and predicted model in cartoon. (**C**) ReLo assay in mammalian cells probing the interaction between KH2F2 and KREH2 dsRBDs. (Left and Center) Confocal microscopy images of cells co-transfected with a cytoplasmic 'prey' domain (GFP-dsRBDs) and a plasma-membrane-targeted 'bait' domain (mCherry-KH2F2). We detect interaction by recruitment of the GFP-labeled prey to the plasma membrane, as seen for dsRBD6 with KH2F2 (top row). The non-interacting pair (bottom row) shows cytoplasmic localization of the GFP-labeled prey. A white line on each image marks the path used for line profile analysis. Scale bars are 5 µm. (Right) ReLo line profiles showing the normalized fluorescence intensity of the GFP (green) and mCherry (magenta) signals along the indicated lines. Co-localization of both peaks indicates an interaction.

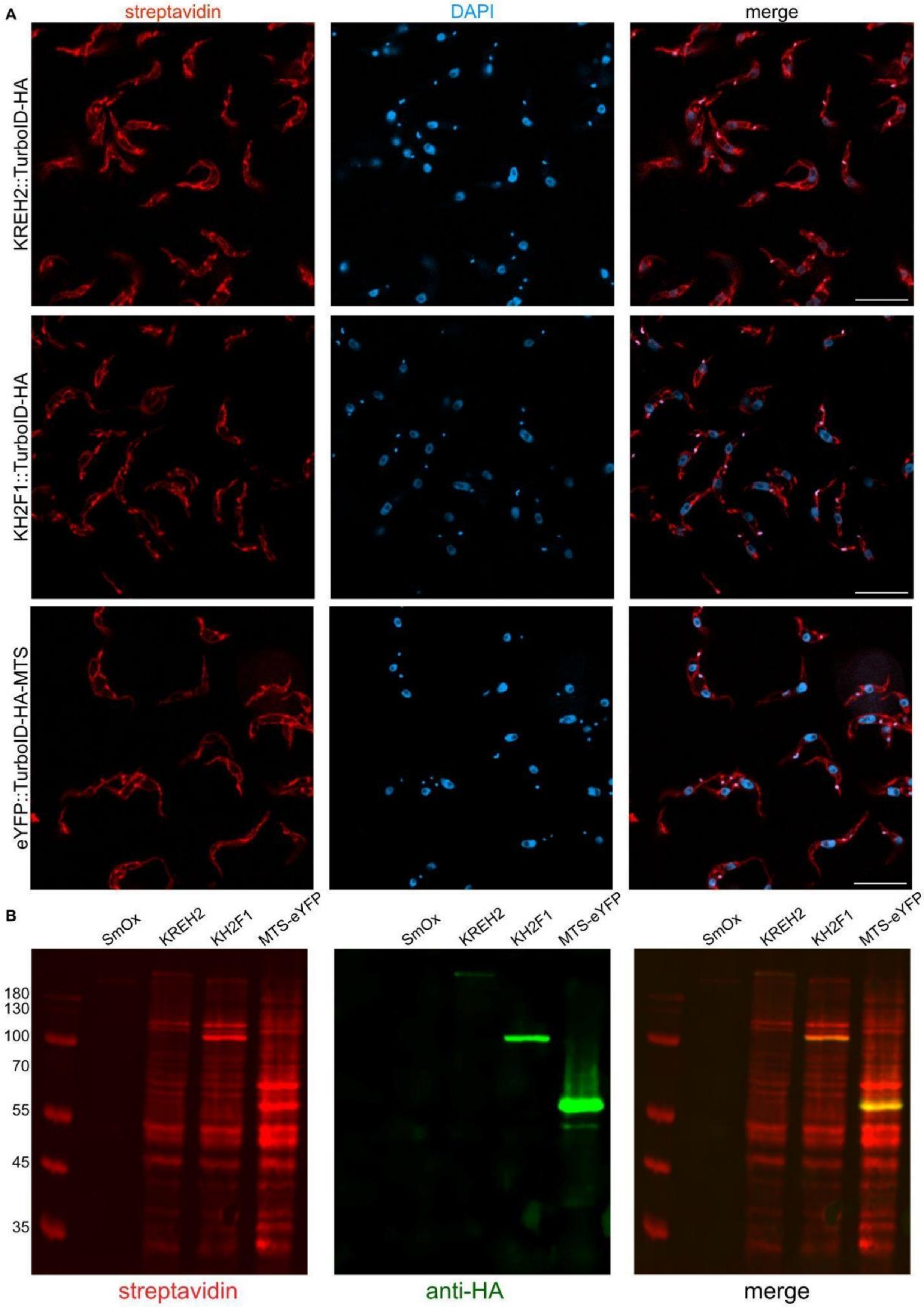
A
streptavidin
DAPI
merge
KREH2::TurboID-HA
KH2F1::TurboID-HA
eYFP::TurboID-HA-MTS
B
SmOx
KREH2
KH2F1
MTS-eYFP
180
130
100
70
55
45
35
streptavidin
anti-HA
merge

**Figure S5: Validation of TurboID cell lines.** (**A**) Fluorescence microscopy images of biotinylation (red) in cells expressing KREH2::TurboID-HA, KH2F1::TurboID-HA, and MTS-TurboID-HA-eYFP. Nuclei were stained with DAPI (cyan). (**B**) Western blot analysis of the biotinylation caused by the TurboID tag (red) and the presence of the HA tag (green) in parental cells (SmOx) and cells expressing KREH2::TurboID-HA, KH2F1::TurboID-HA, and MTS-TurboID-HA-eYFP. The predicted molecular weights of the expressed fused proteins are ~275, 96, and 65 kDa, respectively.

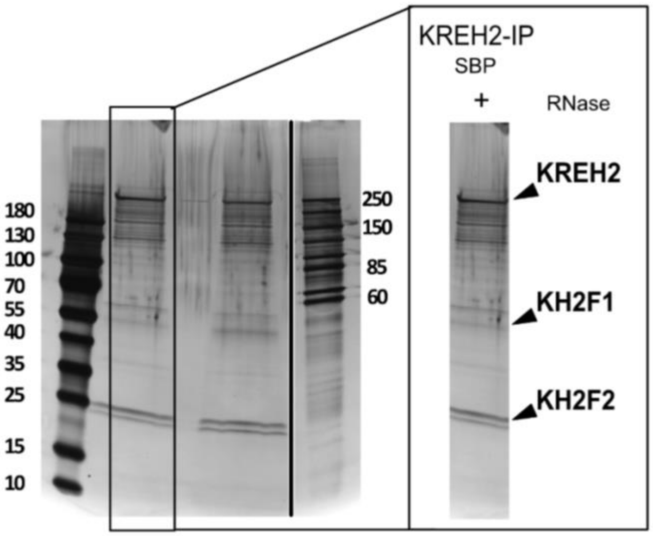


**Figure S6: Immunoprecipitation of endogenous KREH2 tagged with an SBP-3C-ProteinA-tag.** Elution fractions from the streptavidin resin were analyzed by SDS-PAGE on a 4–20% precast gradient gel followed by silver staining. The molecular-weight markers and KREH2 sample shown were derived from non-adjacent lanes of the same gel; intervening lanes containing unrelated samples were omitted, and the spliced regions are indicated by vertical lines. The complete uncropped gel is provided in the Source data.

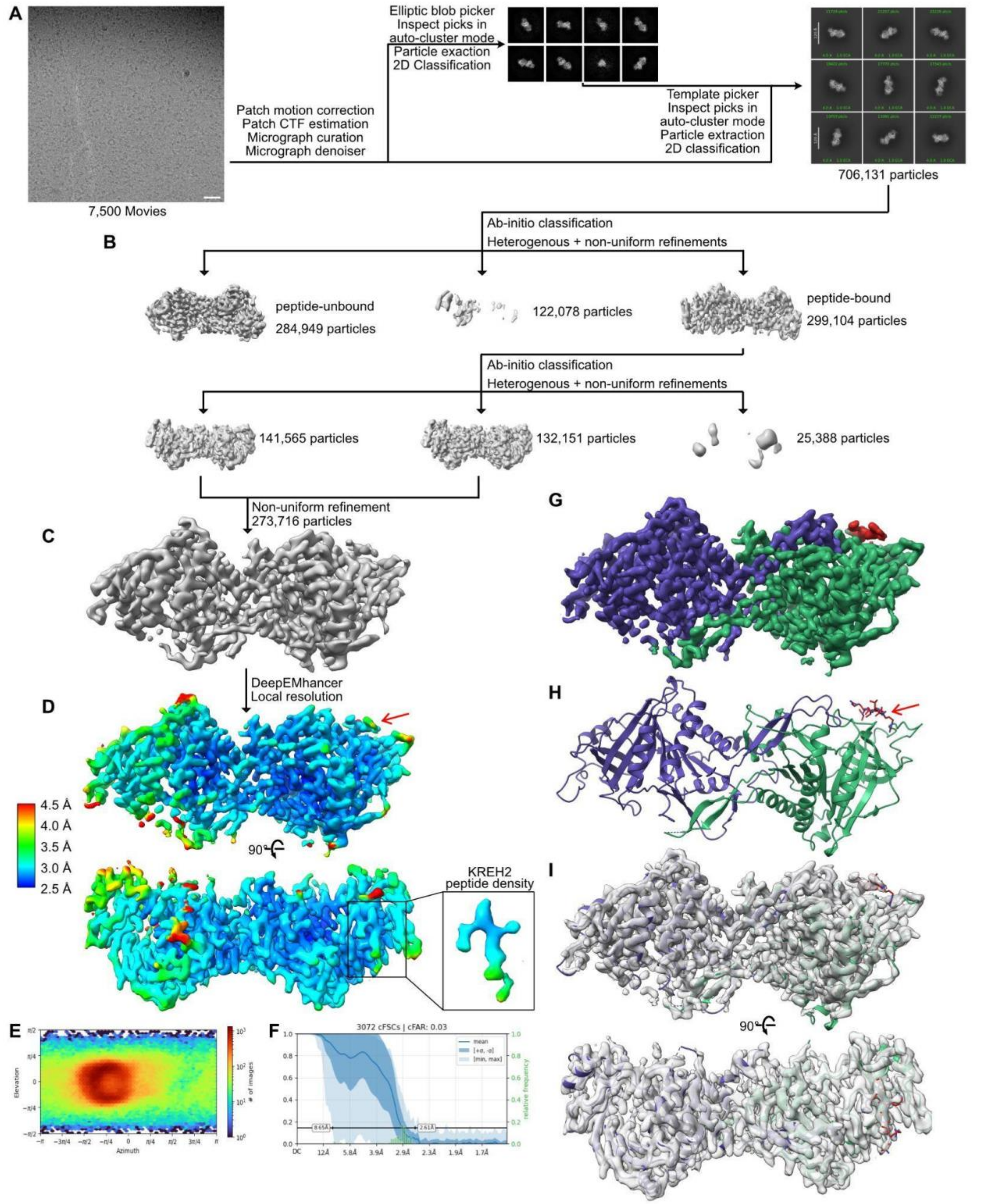


**Figure S7: Cryo-electron microscopy data processing workflow for RESC1-RESC2-KREH2**. (**A**) An exemplary micrograph indicating even particle distribution, with the scale bar corresponding to 100 Å, and pre-processing workflow with motion correction, CTF estimation, particle picking, and 2D classification. (**B**) Particle sorting process by ab initio and heterogeneous refinement. (**C**) Non-uniform refinement of the final particle stack, followed by postprocessing with DeepEMhancer. (**D**) Two orientations of the final map post-processed with DeepEMhancer, colored by local resolution. The red arrow and box indicate the peptide density on the map, with a zoom box showing only the peptide density. (**E**) Angular distribution map of particle orientation of the final non-uniform refinement step done in CryoSPARC. (**F**) Final resolution of the reconstruction, as calculated by CryoSPARC's implementation of the Gold Standard Fourier Shell Correlation, with a resolution of 2.9 Å when adopting a cutoff of 0.143. (**G**) The final post-processed EM map (2.9 Å resolution) of RESC1-RESC2 (dark green, purple) heterodimer and KREH2 peptide (maroon) complex. (**H**) Cartoon representation of the atomic model, with colors corresponding to the map, and a red arrow indicating the KREH2 peptide. (**I**) Two views of RESC1-RESC2 and KREH2 peptide complex atomic model fitted in DeepEMhancer post-processed cryo-EM map.

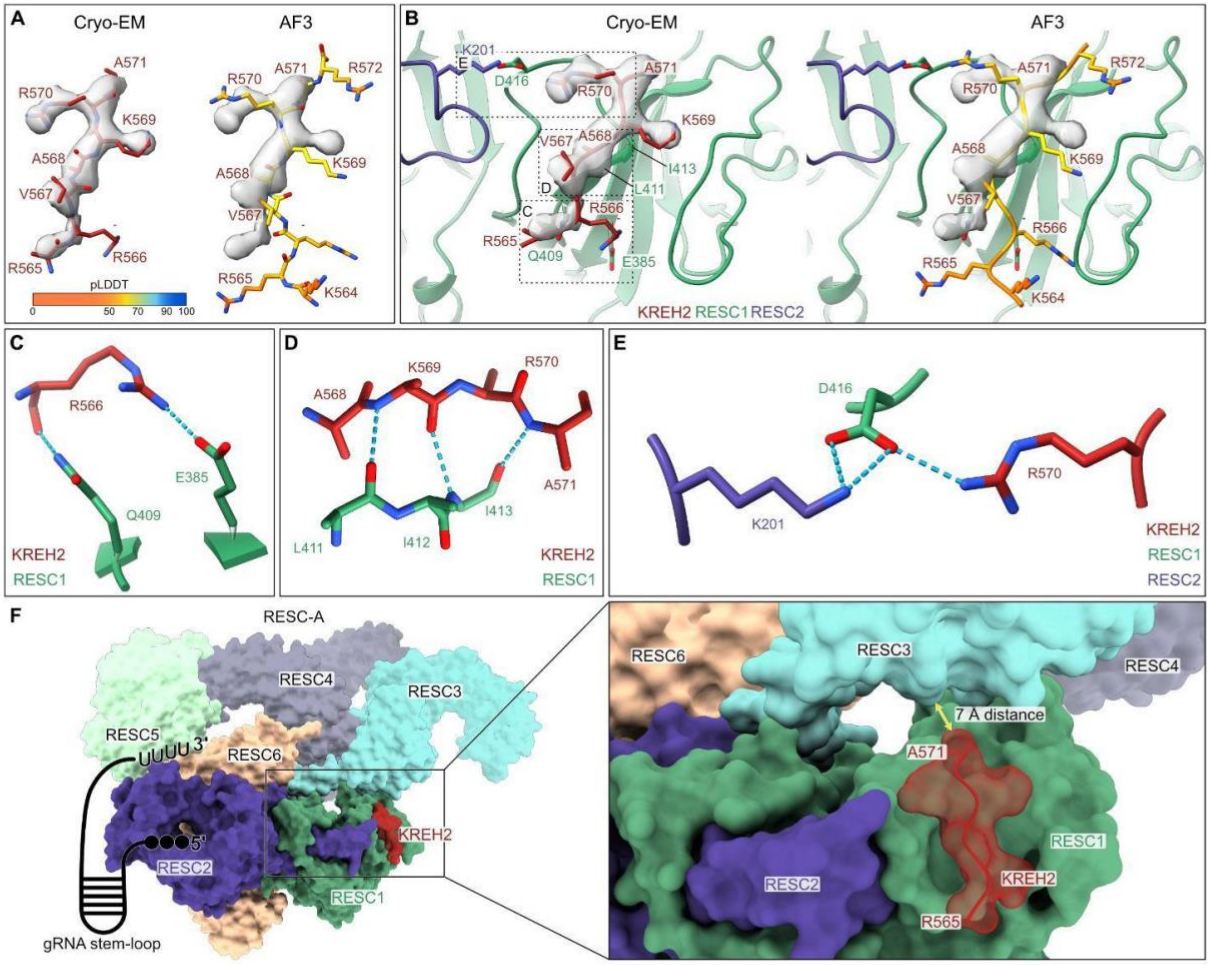


**Figure S8: KREH2 peptide binding to RESC1-RESC2.** (**A**) Zoomed-in view of the density for the KREH2 peptide with the model built in maroon (left) or the AlphaFold3 prediction colored by pLDDT (right). (**B**) Zoomed-in view of the interaction interface between RESC1-RESC2 and the KREH2 peptide, based on modeling into the cryo-EM map (left); the unbiased AlphaFold3 prediction colored by pLDDT for comparison (right). (**C,D,E**) Zoomed-in views of the main contacts between the KREH2 peptide and RESC1-RESC2. Blue dashed lines indicate possible hydrogen bonds. (**F**) Superposition of the cryo-EM structure of RESC1-RESC2-KREH2 with RESC-A (PDB:8FN6). Proteins represented as a surface view, KREH2 peptide as a transparent surface with cartoon. The distance between the last modeled residue of the KREH2 peptide and RESC3 is annotated in yellow.

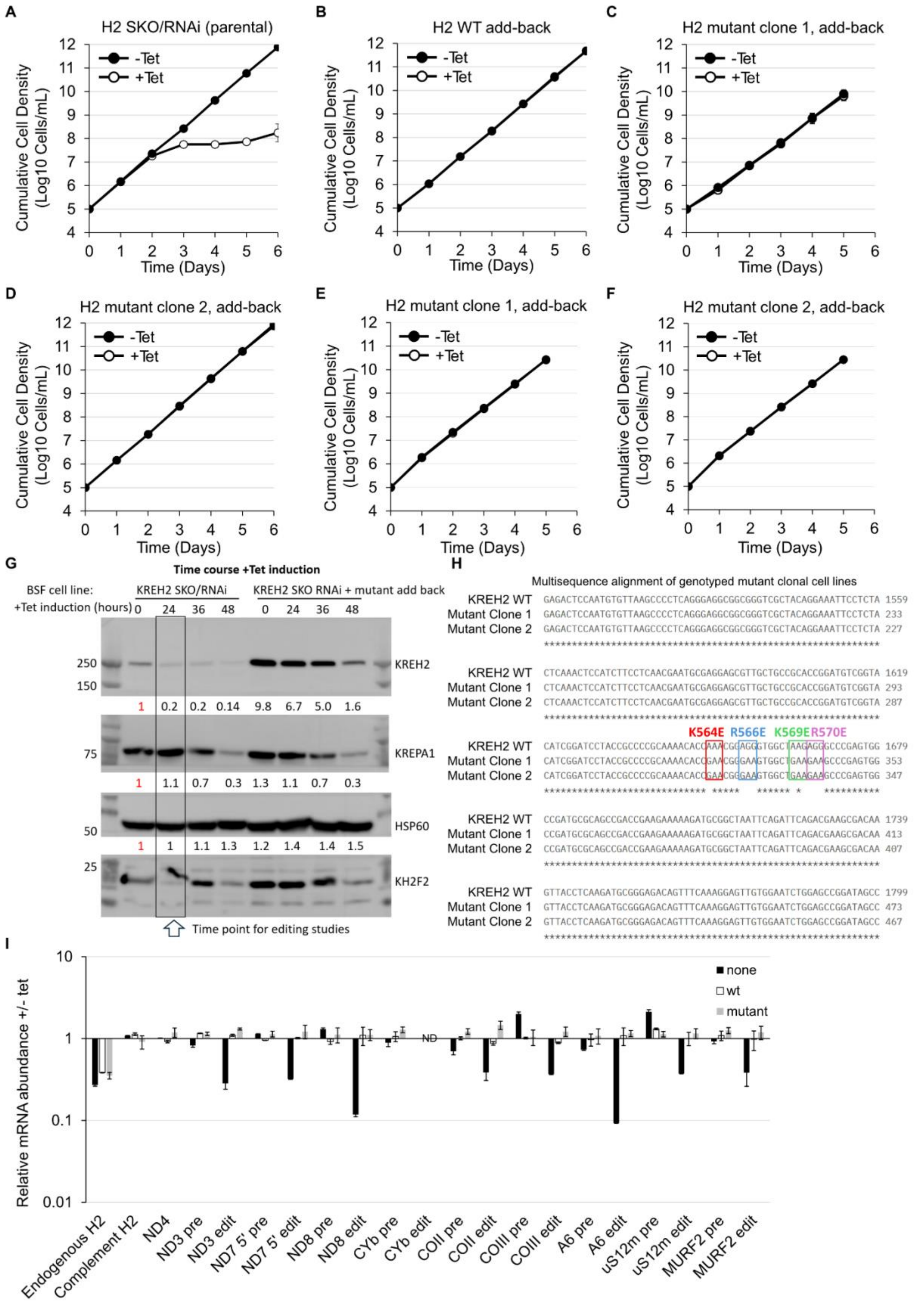
A
H2 SKO/RNAi (parental)
B
H2 WT add-back
C
H2 mutant clone 1, add-back
D
H2 mutant clone 2, add-back
E
H2 mutant clone 1, add-back
F
H2 mutant clone 2, add-back
-Tet
+Tet
Cumulative Cell Density (Log10 Cells/mL)
Time (Days)
G
Time course +Tet induction
BSF cell line:
+Tet induction (hours)
KREH2 SKO/RNAi
KREH2 SKO RNAi + mutant add back
0 24 36 48 0 24 36 48
250
150
KREH2
1 0.2 0.2 0.14 9.8 6.7 5.0 1.6
75
KREPA1
1 1.1 0.7 0.3 1.3 1.1 0.7 0.3
50
HSP60
1 1 1.1 1.3 1.2 1.4 1.4 1.5
25
KH2F2
Time point for editing studies
H
Multisequence alignment of genotyped mutant clonal cell lines
KREH2 WT GAGACTCCAATGTGTTAAGCCCCTCAGGGAGGCGGCGGGTCGCTACAGGAAATTCCTCTA 1559
Mutant Clone 1 GAGACTCCAATGTGTTAAGCCCCTCAGGGAGGCGGCGGGTCGCTACAGGAAATTCCTCTA 233
Mutant Clone 2 GAGACTCCAATGTGTTAAGCCCCTCAGGGAGGCGGCGGGTCGCTACAGGAAATTCCTCTA 227
KREH2 WT CTCAAACTCCATCTTCCTCAACGAATGCGAGGAGCGTTGCTGCCGCACCGGATGTCGGTA 1619
Mutant Clone 1 CTCAAACTCCATCTTCCTCAACGAATGCGAGGAGCGTTGCTGCCGCACCGGATGTCGGTA 293
Mutant Clone 2 CTCAAACTCCATCTTCCTCAACGAATGCGAGGAGCGTTGCTGCCGCACCGGATGTCGGTA 287
K564E R566E K569E R570E
KREH2 WT CATCGGATCCTACCGCCCCGCAAAACACCAAACGGAGGGTGGCTAAGAGGGCCCGAGTGG 1679
Mutant Clone 1 CATCGGATCCTACCGCCCCGCAAAACACCGAACGGGAAGTGGCTGAAGAAGCCCGAGTGG 353
Mutant Clone 2 CATCGGATCCTACCGCCCCGCAAAACACCGAACGGGAAGTGGCTGAAGAAGCCCGAGTGG 347
KREH2 WT CCGATGCGCAGCCGACCGAAGAAAAAGATGCGGCTAATTCAGATTCAGACGAAGCGACAA 1739
Mutant Clone 1 CCGATGCGCAGCCGACCGAAGAAAAAGATGCGGCTAATTCAGATTCAGACGAAGCGACAA 413
Mutant Clone 2 CCGATGCGCAGCCGACCGAAGAAAAAGATGCGGCTAATTCAGATTCAGACGAAGCGACAA 407
KREH2 WT GTTACCTCAAGATGCGGGAGACAGTTTCAAAGGAGTTGTGGAATCTGGAGCCGGATAGCC 1799
Mutant Clone 1 GTTACCTCAAGATGCGGGAGACAGTTTCAAAGGAGTTGTGGAATCTGGAGCCGGATAGCC 473
Mutant Clone 2 GTTACCTCAAGATGCGGGAGACAGTTTCAAAGGAGTTGTGGAATCTGGAGCCGGATAGCC 467
I
Relative mRNA abundance +/- tet
none
wt
mutant
ND
Endogenous H2
Complement H2
ND4
ND3 pre
ND3 edit
ND7 5' pre
ND7 5' edit
ND8 pre
ND8 edit
CYb pre
CYb edit
COII pre
COII edit
COIII pre
COIII edit
A6 pre
A6 edit
uS12m pre
uS12m edit
MURF2 pre
MURF2 edit

**Figure S9: *In vivo* experiments.** (**A-F**) Functional complementation of a BSF KREH2 SKO/RNAi cell line: Parental line (A); wt add-back (B) and two clones of mutant KREH2 add-backs tested independently in two separate labs (C-F). A-D are biological duplicates of technical triplicates. These plots include error bars, which are not apparent in B, C, and D because the readings in these duplicates are very close; (E-F) are single cultures. (**G**) Time course under Tet induction to determine the time for editing studies. (**H**) Multisequence alignment of wild-type and mutant KREH2 variants. Genotyping via Sanger sequencing verified a tetra-substitution K564E, R566E, K569E, and R570E in KREH2 from independent BSF clonal cell lines. (**I**) Consequences to mRNA editing upon KREH2 loss and complementation by constitutively expressed wt or K564E/R566E/K569E/R570E mutant alleles in BSF cells as shown by quantitative RT-PCR analyses. Relative endogenous KREH2, complementary wt or mutant KREH2, never-edited ND4, and pre- and fully-edited mRNA abundances in cells grown for 24 hours in the presence of tet (endogenous KREH2 knockdown via RNAi and exclusive expression of wt or K564E/R566E/K569E/R570E mutant) vs. the absence of tet (endogenous KREH2 expressed) were determined by qRT-PCR. For each target amplicon, the relative change in RNA abundance was determined using the mRNA encoding telomerase reverse transcriptase (TERT) as an internal control. Data are shown as means ± SEM from at least two independent experiments. 'ND' indicates that the mRNAs were not detected in cells grown in both minus tet and plus tet.

**Supplementary Tables**

Due to their large size, supplementary tables can be found in the Supplementary_Tables.xlsx file, with the corresponding titles:

Table S1: Transgenic cell lines generated in this study

Table S2: Primers

Table S3: Plasmids

Table S4: Synthetic gene fragment

Table S5: TurboID MS data

Table S6: BioMark primers (Brenndörfer and Boshart 2010; Carnes et al. 2005, 2018; Meehan et al. 2024)

Table S7: TurboID of KREH2

Table S8: TurboID of KH2F1

Table S9: Co-immunoprecipitation of KREH2

Table S10: Co-immunoprecipitation of RESC2

**Source Data**

Source data for Figures 3B, 4B, 4C, S3, S4, S5, S6 and S9 are provided in the Source_Data.xlsx file.
